# Brain-immune interactions generate pathogen-specific sickness states

**DOI:** 10.64898/2025.12.06.692770

**Authors:** Zuri A. Sullivan, Bertrand J. Wong, Vikrant Kapoor, Paula Kathleen Vincze, Harris S. Kaplan, Pia Giraudet, Brianna R. Watson, Adel Misherghi, Sebastian Lourido, Jeffrey R. Moffitt, Catherine Dulac

## Abstract

In nature, animals encounter diverse pathogens that trigger specific peripheral defense programs and elicit sickness behavior, a set of stereotyped physiological and behavioral changes thought to promote host fitness. Most studies to date have relied on one or a few mouse models of infection, limiting insights into pathogen-specific neuroimmune interactions that generate sickness. We hypothesized that different pathogens might elicit distinct sickness states by engaging different cell types and brain circuits. Using inflammatory models representing bacterial, viral, allergic, parasitic or colitis conditions, we assessed sickness across scales: organismal – behavior and physiology; cellular – brain-wide neural activity; and molecular – single-cell *in situ* transcriptomics in hypothalamus areas associated with social and homeostatic functions affected during sickness. Remarkably, immune challenges elicited unique repertoires of changes across all scales. Our findings reveal pathogen-specific sickness states encoded by the brain across scales, thereby broadening our understanding of how infections make us sick.

## Introduction

The immune and nervous systems enable animals to detect and respond to threats in their environments. The principles underlying neurological and immunological threat detection and response share several common features: peripheral sensing and central processing; dedicated cellular sensors and effectors; the ability to remember past threats and respond more efficiently in the future^1^.

Despite long held assumptions that the brain is insulated from peripheral immune responses, it has become increasingly clear that interactions between the immune and nervous systems are critical for both host defense and homeostasis^1^. For example, tissue resident immune cells in the gastrointestinal tract interact with sensory neurons to efficiently clear bacterial infections^2–5^, and innervation of central lymphoid tissues, such as the spleen, can control the magnitude of immune response^6,7^, highlighting the role of bidirectional communication between the immune and nervous systems. Neurons in the brainstem also regulate the magnitude of peripheral immune responses, and peripheral immune cells traffic to the brain from distal organs, such as the gut and adipose tissue^6,8,9^, illustrating the many brain-body connections underlying neuroimmune interactions. Brain resident immune cells play critical roles in brain development and can be protective or pathogenic in the context of neurodegeneration and aging^10–12^, and immune detection of allergens regulates allergen avoidance^13^. And, critically, neuroimmune interactions underlie sickness behavior, a conserved set of stereotyped changes in physiology and behavior orchestrated by the brain^1^.

Canonical features of sickness behavior include fever, anorexia, lethargy, altered sleep, reduced grooming, and social withdrawal. Recent work has begun to uncover neuronal circuits in the hypothalamus and brainstem that regulate many of these behaviors^14–17^. Much of this work has leveraged an experimental model in which mice injected with lipopolysaccharide (LPS), a component of the cell envelope of gram-negative bacteria, exhibit an acute sickness state that mimics bacterial infection.

LPS-evoked inflammation is mediated by the initial detection of LPS by toll-like receptor 4, a pattern recognition receptor (PRR) that initiates innate immune responses^18^. PRRs detect pathogen-associated molecular patterns (PAMPS), which are conserved structures shared across classes of microbes, enabling rapid detection of infection and initiation of clearance as well as instructing adaptive responses that confer immunological memory^19,20^. Importantly, PRRs are highly diversified in order to enable the detection of the different molecular patterns presented by different pathogens so as to elicit appropriate defense responses^19^. Effector mechanisms that deal with one type of pathogen are not necessarily effective for others. For example, the production of antimicrobial peptides in the gastrointestinal tract helps to clear bacterial pathogens but likely has a minimal impact on viruses or parasites^21^, whereas antiviral responses, mediated in large part by interferon signaling, shut down host cell protein synthesis in order to impede viral propagation^22^. Finally, clearance of macroparasites often involves expulsion rather than direct killing, due to their large size^23,24^. Thus, a modular organization of pathogen detection and response is critical for effective host defenses.

The specificity with which the innate immune system deals with different classes of pathogens raises the question of whether this specificity extends to the neuroimmune interactions that regulate sickness behavior. Indeed, by contrast to the immune system, many of the brain responses to threats appear to be generic rather than specific. Different predator odors engage common neural circuits involving the parabrachial nucleus and periacqueductal gray^25–27^, and sensing of bitter compounds, which typically signal toxicity, does not involve specific discrimination among various bitter ligands^28,29^. In both examples, these neural responses to various threats lead to a common avoidance behavior. Whether the neural response to different pathogens is generic, as in the above examples, or specific, as in the peripheral immune system, is unknown and has important implications for understanding the breadth of molecular and cellular mechanisms that lead to sickness as well as the basis of chronic conditions, such as long COVID, chronic fatigue and other post-acute infection syndromes, which may emerge following an episode of acute infection^30,31^.

In view of the diversity of pathogens that exist in nature, the common and almost sole reliance on LPS as a model of sickness has limited our ability to assess whether different pathogens elicit distinct sickness states. A study showing that antibacterial and antiviral immunity have distinct metabolic needs that directly impact survival supports the hypothesis that different pathogens might elicit distinct sickness states^32^. We sought to test this hypothesis by comparing sickness responses to diverse immunological challenges that represent distinct pathogen types (bacterial, viral and parasitic) and noninfectious inflammatory states (allergy and colitis). We explored sickness across biological scales: at the organismal level, we tested several behavioral and physiological changes; at the cellular level, we looked at neural activity across the entire brain via detection of the immediate-early gene FOS; at the molecular level, we established a single-cell, spatially resolved transcriptomic atlas of the hypothalamic preoptic area, which regulates numerous social and homeostatic behaviors that are altered during sickness. Remarkably, across biological scales, we discovered that neuroimmune interactions generate diverse, pathogen-specific sickness states. Our findings provide new insights into the contribution of sickness to host defense and open new avenues of research into mechanisms underlying the brain responses to infection and inflammatory states.

## Results

### Different immunological challenges have distinct effects on animal physiology

We used distinct sterile models to elicit immune responses that mirror diverse pathogens and non-infectious immune challenges. These included LPS, a molecule derived from the cell wall of gram-negative bacteria, which models bacterial infection; polyinosinic:polycytidylic acid (pI:C), a synthetic mimic of double-stranded RNA which models viral infection^33,34^; soluble Toxoplasma antigen (STAg) isolated from *Toxoplasma gondii*, which models parasite infection^35–37^; bee venom phospholipase A2, the primary allergic component of bee venom (PLA2), which models acute allergic immunity^38^; and dextran sodium sulfate (DSS), a chemically-induced model of colitis^39^ (Fig. 1A). With the exception of DSS, which is administered in the drinking water over several days, all compounds were administered via intraperitoneal injection and doses were optimized to elicit acute sickness without severe morbidity^15,38,40^. As expected, each model induced a distinct repertoire of circulating cytokines (Fig. 1B, Fig. S1C), and a classifier trained on the serum concentration of all (Fig. S1A) or a subset (Fig. S1B) of these cytokines predicts the identity of the sickness mode. This is consistent with the notion that peripheral responses vary across different immune challenges. LPS-injected mice showed increases in numerous cytokines involved in antibacterial immunity, pI:C-injected mice showed increases in type-1 interferon – a master regulator of antiviral immunity – and PLA2-injected mice showed increases in cytokines involved in type-2 immunity: interleukin- (IL) 4, IL-5, and IL-13^22,41^. DSS-challenged mice showed changes in a subset of type-1 cytokines (IL-12p70 and IL-1b)^39^, and STAg had minimal impact on peripheral cytokines, but, along with DSS, elicited elevated levels of IL-7 and IL-15 (Fig1B).

**Figure 1.**
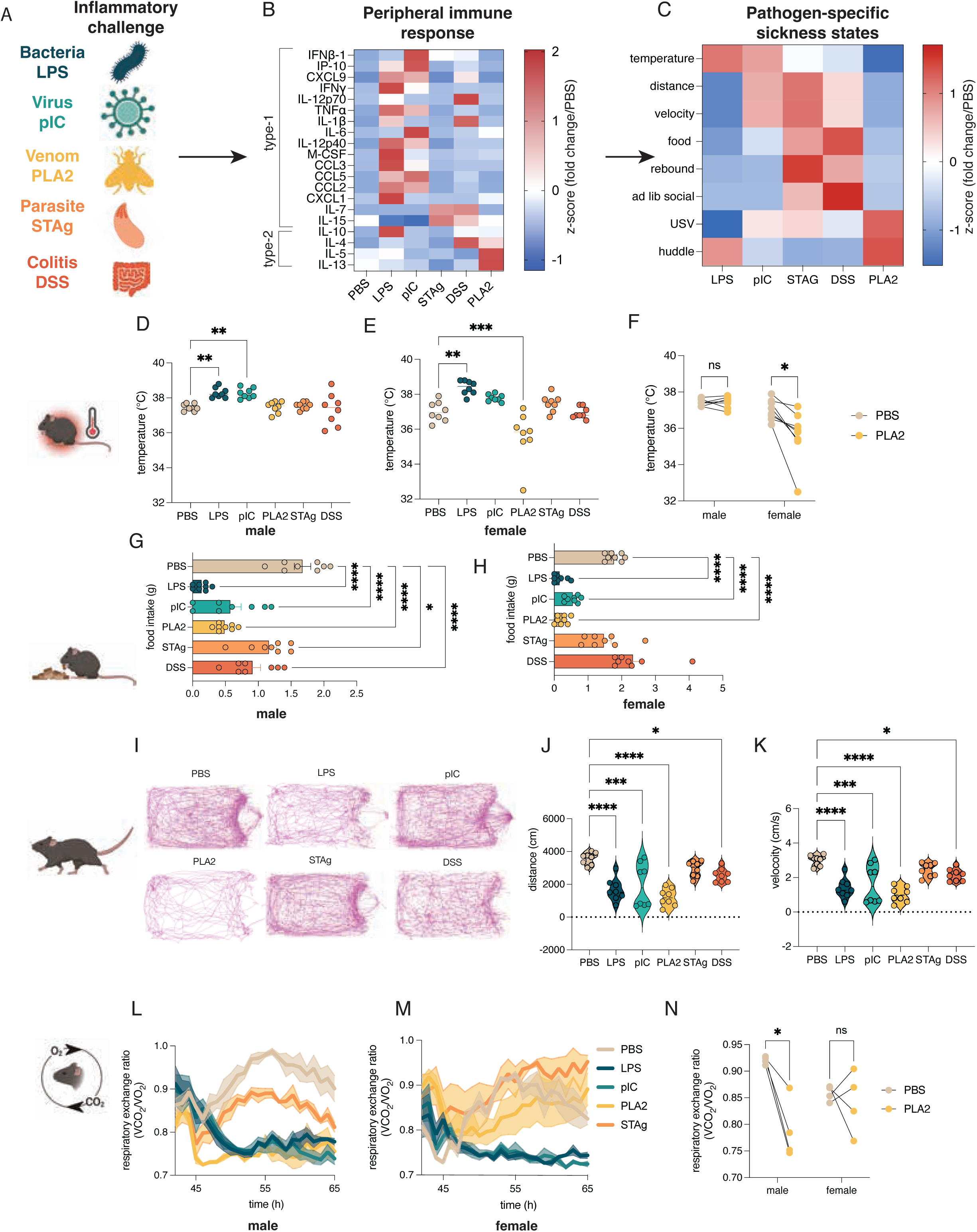
Systemic physiologic changes across sickness states. (A) Models used to elicit pathogen-specific sickness states (B) Serum cytokines induced in response to different sickness models measured by multiplexed enzyme linked immunosorbent assay (ELISA). (C) Summary of physiologic and behavioral changes elicited by different sickness models, summarizing data in Fig.1, Fig.S1, Fig. 2, Fig. S2. (D) Body temperature during different sickness states in males (E) Body temperature during different sickness states in females (F) Hypothermia in PLA2-injected mice in females versus males (G) Food intake over 4h during different sickness states in males (H) Food intake over 4h during different sickness states in females (I) example traces of locomotor activity over 10 minutes during different sickness states (J) distance traveled and (K) average velocity during different sickness states in males and females combined (L) respiratory exchange ratio (RER) during different sickness states in males, quantified by indirect calorimetry (M) RER during different sickness states in females, quantified by indirect calorimetry (N) comparison of RER in male vs. female mice after PLA2 injection. Heat maps represent z-scored data. p<0.05* p<0.01** p<0.001*** p<0.0001**** based on ordinary one-way ANOVA with Dunnett’s multiple comparisons test. All measurements taken at 4h post-injection (acute models) or day 3 (DSS).

To assess whether the five models of immune challenges induced different organismal phenotypes, we initially measured canonical features of sickness, including body temperature, food intake, and locomotion (Fig. 1C). All experiments were performed in C57BL/6 mice and a subset was additionally done with FVB mice due to known strain differences in social behavior^42^. As previously described, both LPS and pI:C elicited a fever response in C57BL/B6 male mice^15^, and LPS in C57BL/B6 female mice. In FVB mice, both males and females exhibited a febrile response to LPS and pI:C. In contrast, neither STAg nor DSS induced a change in temperature. and PLA2 induced hypothermia in C57BL/B6 females, and in both sexes on the FVB background (Fig. 1D-F, C57BL/6; Fig S1D&E, FVB). This is consistent with the hypothermic response observed during anaphylaxis and the well-described sexual dimorphism seen in allergic immunity^43,44^. We next tested how these different models influenced food intake. In C57BL/6 males, all models caused a reduction in food intake, while in C57BL/B6 females, only LPS, pI:C, and PLA2 reduced food intake. In FVB mice, food intake was reduced in LPS-, pI:C, and PLA-2 challenged males, in LPS-, pI:C-, PLA2-, and STAg-injected females, and was increased in DSS-challenged females, during the four hours of observation (Fig. 1G&H; Fig. S1 F&G). Measuring locomotion, we found that all models, except STAg, elicited a reduction in average velocity and distanced traveled in C57BL/B6 mice, and all five models caused reduced locomotion in FVB mice during a 10-minute observation period 4 hours after injection. However, the reduced locomotion was greater in LPS- and PLA2challenged mice than in other models (Fig. 1I-K; Fig. S1H&I).

Based on the previously reported significance of alterations in systemic metabolism for survival during different types of infection^32^, we next assessed the respiratory exchange ratio (RER) using a continuous laboratory animal monitoring system (CLAMS). RER is quantified as the ratio of carbon dioxide production to oxygen consumption, and is used as a proxy for fuel utilization, with an RER of 1.0 indicating that an animal is relying on glucose, and an RER of 0.7 indicating a reliance on fatty acids. Over a 24-hour recording period, we found marked differences in RER across our models. Of note, mice were allowed to acclimatize in CLAMs devices for 24 hours and were challenged with given inflammatory reagents at approximately ZT12, when mice are active. This initial challenge caused an acute drop in RER in all conditions, including saline, likely due to the stress of handling and intraperitoneal injection. In males, LPS, pIC, and PLA2 models resulted in a significant drop in RER upon injection that persisted for the duration of the recording, whereas STAg (Fig.1L, Fig. S1J&K) and DSS (Fig. S1N) had no significant impact on RER in males. In females, LPS and pIC had profound impacts on RER, whereas PLA2, STAg (Fig. 1M,N; Fig. S1L, M) and DSS (Fig. S1O) had no effect. As in body temperature measurements, LPS and pI:C were more similar in their effects than PLA2, STAg, and DSS with sex being an important variable in determining the metabolic response to PLA2 (Fig. 1N). Together, these results indicate that different immunological challenges elicit distinct core physiological responses including food intake, body temperature, locomotion, and metabolism.

LPS and pI:IC both represent “type-1” immune challenges, which elicit distinct but overlapping repertoires of circulating cytokines, whereas PLA2 induced a distinct, type-2 cytokine signature, DSS induced a subset of type-1 cytokines, and both STAg and DSS induced production of circulating IL-7 and IL-15 (Fig 1, Fig. S1C). The relative similarity of peripheral responses to LPS and pI:IC compared to the other tested models may help explain why these two challenges elicit similar physiological responses during sickness. Across experiments, STAg and DSS were also generally more similar to each other than to the other models, and PLA2 was distinct from other models by some metrics (temperature, cytokines, metabolism) and similar by others (locomotion, food intake). Together, these results demonstrate that different immunological challenges elicit distinct sickness states, in which the brain orchestrates changes in homeostatic systems in response to the detection of pathogen-associated signals and that the impact of neuroimmune interactions on these states may slightly differs across sexes.

### Different immune challenges distinctly affect social behavior

Sickness has been shown to trigger social withdrawal, which helps protect uninfected animals from social transmission of infection^1,14^. We recently characterized distinct hypothalamic cell populations that regulate the need and the satiety for social interaction, respectively^42^. Neurons in the medial preoptic nucleus (MPN) characterized by coexpression of *Mc4r* and *vGlut2* (MPN^vGlut2+ Mc4r+^) are specifically activated by social isolation and play a causal role in promoting social drive and eliciting a rebound in social interaction upon reunion. A distinct MPN cell type, marked by *vGat* and *Trhr* expression (MPN^vGat+Thrh+^), is activated during social rebound and signals the satiation of social need^42^.

We used two different experimental paradigms to assess how various sickness models influence social interactions and the activity of the corresponding neural circuit. In an initial assay, we tested how sickness influences *ad libitum* social interactions by allowing socially housed female FVB mice, identified in our previous study as highly sensitive to social isolation^42^, to interact with a littermate in a recording arena for 10 minutes. As predicted, saline-injected mice showed minimal social interaction with a familiar mouse, preferring to explore the arena. LPS-, pI:C, and PLA2-injected animals also exhibited minimal engagement in social behavior (Fig. 2A,B). In contrast, STAg- and DSS-challenged animals showed significantly higher levels of social interaction than did saline controls, indicating that these conditions drive pro-social behavior independent of social isolation (Fig. 2A,B). Strikingly, STAg and DSS elicited *fos* activity in MPN^Mc4r+^ isolation neurons suggesting enhanced social drive. Indeed, the extent of *fos* induction in STAg and DSS treated animals was comparable to that observed in saline-injected mice during isolation (Fig. 2C). Together, these results demonstrate that STAg and DSS conditions contrast with other sickness models in promoting prosocial behavior in the absence of isolation and induce activity in neurons that promote social interaction.

**Figure 2.**
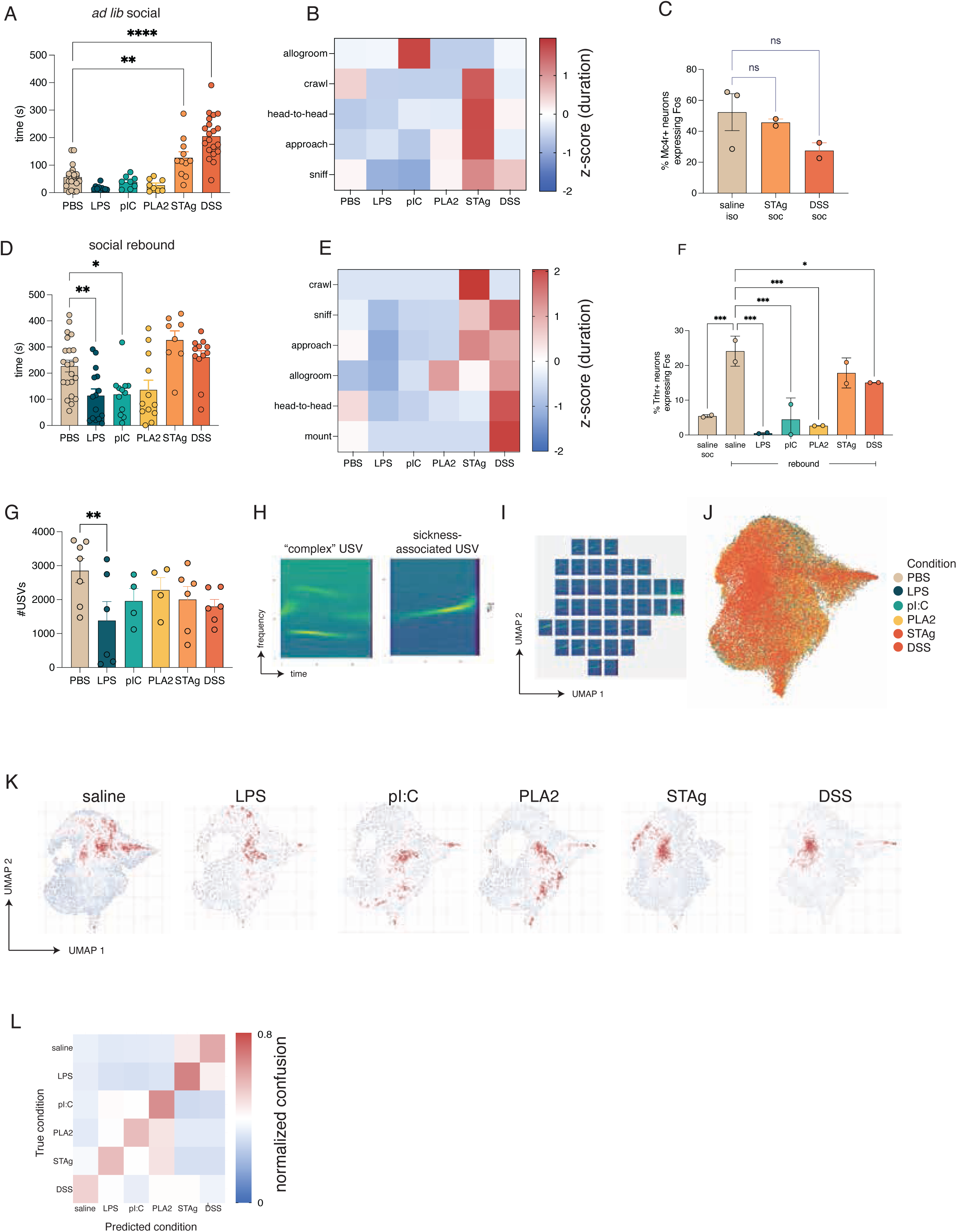
Altered social behavior during different sickness states. (A) Duration of *ad libitum* social interaction in FVB female mice recorded over 10 minutes. (B) Individual social behaviors quantified in (A). (C) *Fos* expression in *Mc4r*+ ‘isolation’ neurons in isolated saline-injected mice or socially housed STAg- or DSS-challenged mice (D) Social interaction in FVB females recorded over 10 minutes following 3 days of social isolation. (E) Individual social behaviors quantified in (D). (F) *Fos* expression in *Trhr*+ ‘reunion’ neurons during social rebound. (G) Number of ultrasound vocalizations (USVs) elicited during social rebound. (H) Representative spectrograms of USVs elicited during social rebound in saline-injected or sick mice. (I) UMAP coordinates of representative spectrograms (J) UMAP projection of all USVs, colored by condition (K) UMAP projection of USV spectrogram data separated by condition, colored by density (L) Normalized confusion matrix for random forest classifier trained on USV acoustic features quantified using WarbleR. Heat maps in (B) and (E) represent z-scored data. p<0.05* p<0.01** p<0.001*** p<0.0001**** based on ordinary one-way ANOVA with Dunnett’s multiple comparisons test. All measurements taken at 4h post-injection (acute models) or day 3 (DSS).

Next, we tested how different sickness models influence social rebound behavior in response to isolation and the activity of the corresponding neuronal populations. FVB female mice were isolated from their cagemates for 3 days and monitored in a behavioral arena during reunion with a cagemate. As expected, saline-injected mice showed high levels of social interaction after isolation (Fig. 2D). In contrast, LPS- and pI:C, animals showed minimal social interaction after isolation, while PLA2 showed a trend towards reduced sociality, suggesting that these sickness models interfere with the normal increase in social drive during isolation (Fig. 2D&E). In addition to the range of social behaviors observed previously (sniff, approach crawl, head-to-head contact, allogrooming)^42^, we also observed mounting behavior in DSS-challenged FVB females and PLA2-challenged C57BL/6 females during social rebound (Fig. 2E, Fig. S2D). As in the previous assay, STAg- and DSS-challenged animals showed high levels of social interaction, confirming an increase in prosocial behavior independent of social isolation. Accordingly, we found that saline-, STAg-, and DSS-treated animals showed a trend towards higher levels of *fos* expression in MPN^Trhr+^ neurons, which encode social satiety during rebound, while LPS, pI:C-, and PLA2-challenged animals showed minimal *fos* expression after social interaction (Fig. 2F). Together, these results indicate that some sickness models (LPS, pI:C, and PLA2) inhibit social interaction and the activity of social homeostasis circuits, whereas other sickness models (STAg, DSS) promote social interactions and enhance the activity of circuits underlying social drive.

Similar experiments were performed in C57BL/6 animals, which typically showed a markedly reduced sensitivity to social isolation compared to FVB animals^42^. STAg and DSS challenge resulted in a modest, but not statistically significant, increase in *ad libitum* social interaction in C57BL/6 mice, while pI:C and PLA2 resulted in highly variable responses (Fig. S2A&B). Because these mice showed minimal social interaction before and after isolation, we were unable to detect an additional reduction in social interaction after LPS, pI:C, or PLA2 challenge (Fig. S2C&D).

Finally, we measured the production of ultrasonic vocalizations (USVs), a non-locomotor social behavior that increases during social isolation^42^. We found that LPS-injected animals showed a significant decrease in USV production with a nonsignificant trend toward decreased vocalizations in the other models (Fig. 2G). We also noticed distinct patterns in the spectrograms of sick versus healthy mice (Fig. 2H), which led us to explore in more detail the acoustic features of USVs across sickness models as well as the qualitative features of the spectrograms themselves. When the spectrograms emitted across all conditions were projected onto UMAP space, each sickness model emerged as eliciting distinct and overlapping patterns of USV types (Fig. 2I-K). Certain acoustic features of the USVs representing “complexity,” such as entropy and range of dominant frequencies, were significantly altered in LPS, pIC, and PLA2, but not STAg and DSS (Fig. S2E-J). Strikingly, a random forest classifier trained on 35 acoustic features of USVs was able to predict the specific sickness state with more than twice the accuracy from shuffled data (F=0.475 +/- 0.006 vs. 0.166 +/- 0.005), underscoring the unique features of animal vocalizations across various immune challenges (Fig. 2L).

Together, these data demonstrate that different immunological challenges elicit distinct repertoires of social behavior. Remarkably, changes in social homeostasis observed during different immune challenges followed similar patterns as changes in other homeostatic programs. The effects of STAg and DSS were more similar to each other than to other sickness models, as were the effects of LPS and pI:C. This indicates that, while each sickness model elicits a distinct repertoire of behavioral and physiological changes, there are also overlaps between the behavioral and homeostatic changes elicited by certain immune challenges. To a lesser extent, we observed sex-and strain-dependent changes in certain sickness behaviors, suggesting that sickness, like other aspects of the immune response, can be influenced by sex and genetic background^44,45^.

### Brain-wide neural activity patterns differ across diverse immune challenges

We next investigated how different sickness states are encoded in the brain. Based on the different behavioral responses to various immune challenges, we hypothesized that each immune challenge likely triggers a unique pattern of brain activity. To test this idea, we took an unbiased approach and quantified the expression of FOS across the whole brain. FOS is a transcription factor and the product of *fos*, an immediate-early gene with expression rapidly induced upon neuronal excitation, making the quantification of FOS expression in neurons a widely used tool as a proxy for neuronal activation^46^. We performed tissue clearing and whole-mount immunohistochemical staining for FOS, followed by lightsheet imaging to generate 3-dimensional images of FOS across the entire brain 4 hours after each immune challenge, or on day 3 of DSS treatment. We then applied a machine learning model to register these 3-D images to the Allen Brain Atlas and quantify FOS expression within each brain area annotated in the Atlas (Fig. 3A), generating a matrix of 48 animals x 641 brain areas of FOS intensities and a library of images used for downstream analyses. Given the richness and complexity of our dataset, we initially focused on key areas of interest based on the behavioral and physiological changes described earlier with the expectation that future analyses will further delineate the brain-wide significance of the observed activity patterns.

**Figure 3.**
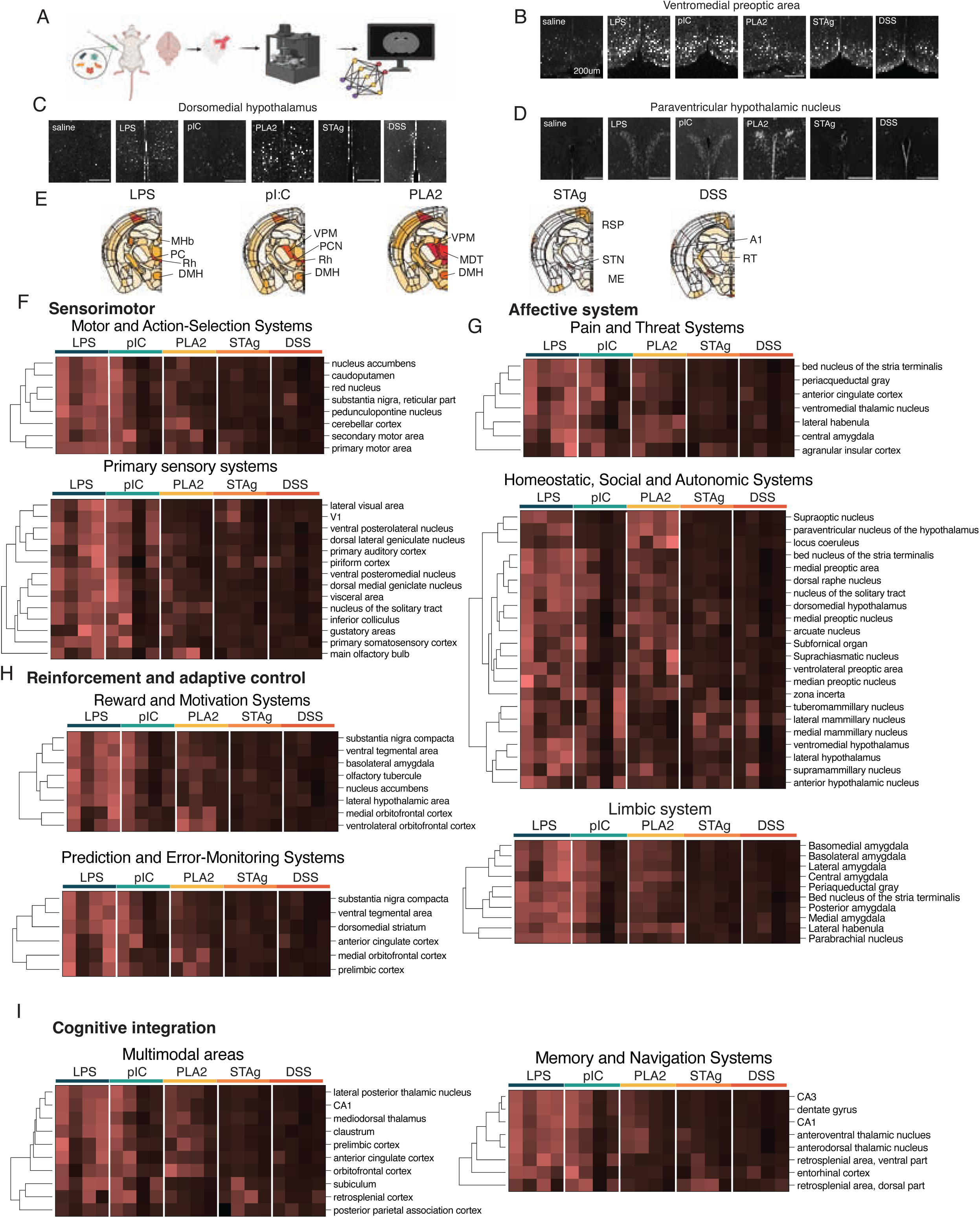
Whole-brain FOS analysis during different sickness states. (A) Pipeline for whole-brain FOS data collection. (B-D) representative images of FOS antibody staining in selected hypothalamic areas. Scale bar = 200um. (E) Graphical heat map of average FOS intensity across coronal sections in all sickness states, normalized to saline controls. (F-I) Hierarchical clustering of FOS intensity normalized to saline control across brain areas grouped by function. Acronyms of brain areas shown in (E) listed in Table S1.

In line with previous work identifying VMPO^LPS^ neurons as key regulators of fever, warmth seeking and appetite during sickness, we confirmed an increase in FOS expression in the ventromedial preoptic area (VMPO) in mice injected with LPS (Fig. 3B). In the dorsomedial hypothalamus (DMH), another thermoregulatory area, we observed increased FOS in LPS- and pI:C-injected animals, which showed a febrile response, and PLA2-injected animals, which showed hypothermia. (Fig. 1D-F; Fig. 3C) Furthermore, we saw strong FOS induction in the paraventricular hypothalamus (PVN) in LPS-, pI:C-, and PLA2-injected animals, suggesting a possible role of the neuroendocrine system in mediating the social and physiological changes we observed in these acute models of sickness (Fig. 3D).

After assessing activation in expected brain areas, we systematically assessed brainwide FOS induction by quantifying FOS intensity in each brain area in our dataset and normalizing to saline controls that were imaged in the same session to get a ratio of FOS in sick vs. healthy animals. The maps of average FOS intensity across coronal sections in each sickness state provided a display of activity in individual areas of interest and, in some cases, could be correlated with observed behavior changes (Fig. 3E, Fig. S3A). For example, the medial habenula, rhomboid nucleus, and bed nucleus of the stria terminals, all involved in anxiety-like behavior, were seen altered in LPS-, pI:C-, and PLA2challenged animals – conditions associated with social withdrawal.

Next, we performed a hierarchical clustering of induced FOS intensity in groups of brain areas related to different biological functions (Fig. 3F-I, FigS3B-F, Table S1). Among brain areas involved in sensorimotor function, we compared FOS in areas related to motor and action selection as well as primary sensory systems. Within motor and action selection systems, including primary and secondary motor cortices, we saw the highest degree of FOS in LPS-injected mice, with intermediate levels in pI:C- and PLA2-injected mice (Fig. 3F), all conditions in which we saw marked reduction in locomotion (Fig. 1I-K). Neurons in the caudal pontine reticular nucleus (RPO), also involved in motor function, showed increased FOS in STAG- and DSS-challenged animals (Fig. S3A).

In primary sensory systems, we saw the greatest increase in FOS in LPS- and pI:Cinjected animals (Fig. 3F). In visual areas, including V1 and the lateral visual cortex, we also saw modest FOS induction in STAg-challenged animals, and in olfactory areas, including the main olfactory bulb and piriform cortex, we saw strong FOS induction in PLA-2- and LPS-challenged animals, as well as after STAg challenge (Fig.3F).

Among brain areas involved in homeostatic and autonomic control, which regulate many of the sickness behaviors we assessed in early experiments, many showed strong FOS induction in the three acute models (LPS, pI:C, and PLA2) that had the strongest impact on feeding, temperature, metabolism, and locomotion, supporting the involvement of areas that regulate these behaviors (Fig. 3G). One exception to this pattern was the mammillary body, involved in learning and memory, in which STAg- and DSS-challenged animals exhibited increased FOS as compared to other hypothalamic areas (Fig. 3G) Other clusters of nuclei were activated across various combinations of immune challenges. Neurons in the lateral hypothalamic area (LH) and arcuate nucleus of the hypothalamus, which regulate feeding behavior, were active in LPS- pI:C-, and PLA2injected animals, all of which displayed reductions in food intake (Fig. 1G&H, Fig 3G). The tuberomammillary nucleus (TMN), which contains histaminergic neurons that regulate sleep-wake cycle, showed increased FOS in all conditions except PLA2 (Fig 3G). The subfornical organ (SFO), a circumventricular organ (CVO) that interfaces with the periphery via a fenestrated blood brain barrier, was robustly activated in LPS- and PLA2- injected animals, and more variably in pI:C (Fig. 3G, Fig.S3A).

Brain areas involved in pain and threat sensing were also highly activated in these conditions, which may be consistent with the reduction in social behavior observed in these animals (Fig. 2). Numerous threat sensing areas, including the bed nucleus of the stria terminalis, periacqueductal gray, central amygdala, and lateral habenula, showed marked increases in FOS intensity in the three models that were associated with reduced social interaction – LPS, pI:C, and pLA2 (Fig. 3G). In addition to these areas, high FOS intensity in the PVN and numerous brain areas in the hypothalamic preoptic area and limbic system (Fig. 3A&G, Fig. S3A,C,D) across these models indicates strong activation of numerous areas involved in social interaction, one of the strongest behavioral phenotypes we observed (Fig. 2).

We next looked at brain areas involved in reinforcement and adaptive control, including systems for reward and motivation, as well as those for prediction and error-monitoring. Here we saw clusters of brain areas (medial orbitofrontal cortex, prelimbic cortex, and ventrolateral orbitofrontal cortex) that were most strongly activated in LPS- and PLA2- injected animals (Fig. 3H). Other brain areas in these categories, particularly those involved in reward, such as the ventral tegmental area, substantia nigra, and nucleus accumbens, showed increased FOS in LPS-injected animals and an intermediate level of FOS induction in PLA2- and pI:C-injected animals (Fig. 3H).

Finally, brain areas involved in higher-order functions showed slightly different patterns of activity. We observed activation of the subiculum, retrosplenial cortex, and posterior parietal association cortex in STAg- and LPS-injected animals. Other cortical areas followed a more similar pattern to subcortical areas, with LPS-, pI:C, and PLA2-injected mice showing the highest FOS activity (Fig. 3I).

A pattern that emerged across several functional categories was apparent sex differences in FOS induction in PLA2-challenged animals, and, to a lesser extent, in pI:C-injected animals, whereas LPS-injected females appeared to show greater inter-individual variability than LPS-injected males (Fig. S3B-F). Together with our physiology data (Fig. 1D-N) these observations suggest that certain sickness states may exhibit more sexual dimorphism than others. We also observed that, in general, LPS elicited stronger patterns of FOS induction than other models, despite significant behavior and physiologic changes observed in all cases. Future work building upon these proof-of-concept studies may perform detailed characterization of individual models, coupling behavior to real-time neural activity recording in targeted brain areas to better link activity to behavior with greater temporal resolution. Overall, these results support the hypothesis that diverse sickness models impact behavior and physiology via coordinated effects on whole-brain activity across functional modalities. Moreover, these data lay the groundwork for a further analysis of the causal relationships between patterns of activity and observed sickness phenotypes, and to understand how inter-individual heterogeneity observed in these responses may contribute to, or impair, the induction of protective physiological states, as reported by others^47,48^.

### Brain-wide neural signatures of distinct sickness states

Following this candidate-based approach, we implemented an unbiased quantitative pipeline to compare brain-wide FOS induction across different sickness states This approach was motivated by the hypothesis that discrete sickness states are characterized by unique cellular FOS expression patterns capable of serving as biomarkers for their subsequent classification. As a preparatory step, the FOS intensity matrix (encompassing 641 discrete brain areas) was subjected to data normalization using a Yeo-Johnson transformation (Fig. 4A). Next, using an adapted Boruta feature selection method, we identified a reduced subset of brain regions (Fig. 4A) that significantly improved clustering of sickness states, as visualized with t-SNE (Fig. 4B, Table S2&3). This reduction enhanced the biological signal by filtering out irrelevant features, improving clustering metrics and separation between conditions. Quantitative validation was performed by calculating two key clustering metrics: the silhouette distance, which measures the cohesion and separation of the clusters, and the Calinski-Harabasz index (also known as the Variance Ratio Criterion), which quantifies the ratio of between-cluster variance to within-cluster variance. Both metrics demonstrated that the feature set identified by the Boruta feature selection improved the resulting data partition (Fig. S4A).

**Figure 4.**
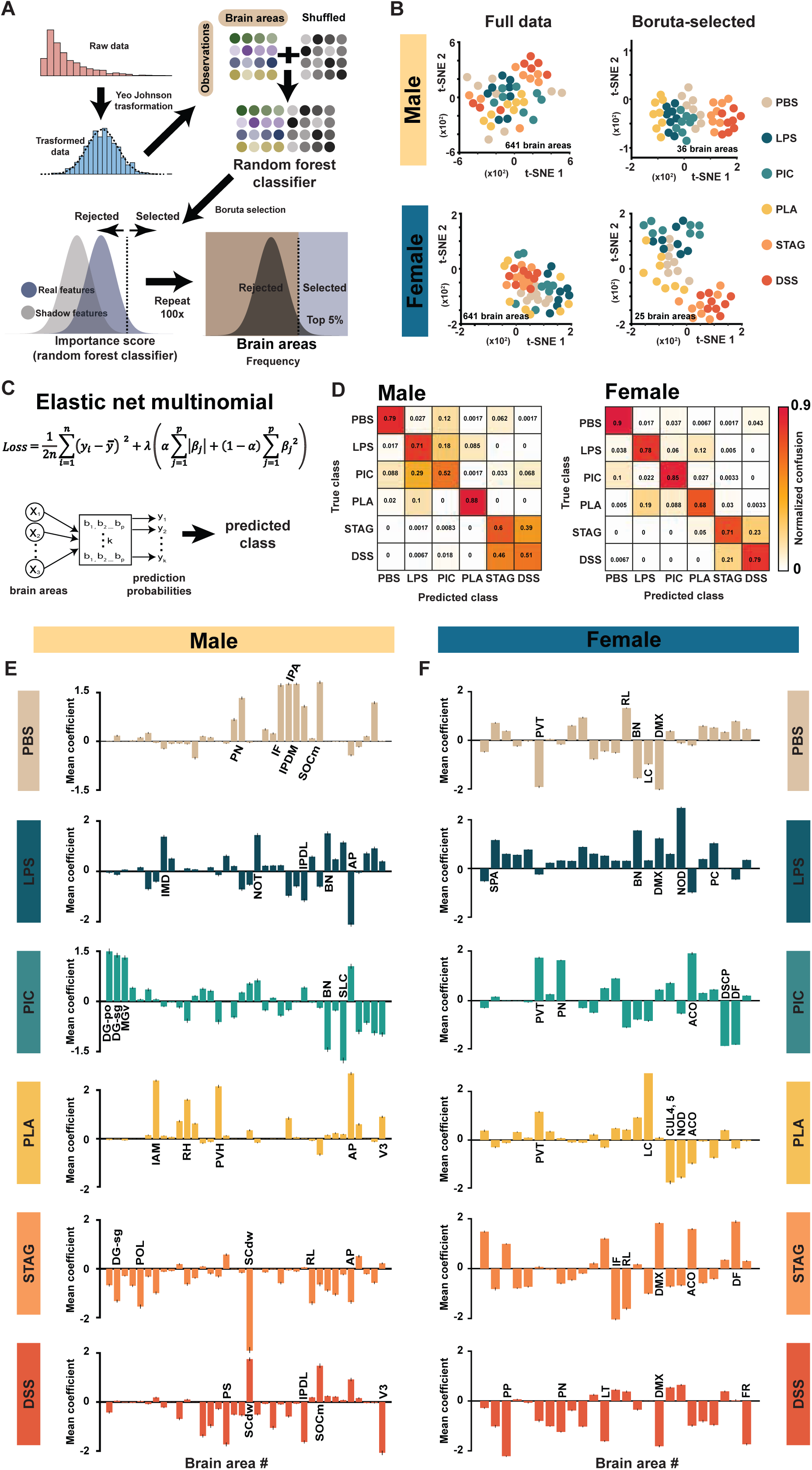
Unbiased quantitative analysis reveals distinct neural signatures for sickness states. (A) Quantitative pipeline for FOS expression data: raw FOS intensity data from 641 brain areas was normalized using a Yeo-Johnson transformation. An adapted Boruta feature selection method was then implemented to identify the most informative brain regions. The process was repeated 100 times, selecting the top 5% of features based on frequency. The final selection yielded 36 key brain regions in males and 25 in females. This approach minimizes bias and prevents overfitting. (B) Clustering Visualization of FOS Data: t-SNE visualization of FOS expression data before (641 areas) and after (36 or 25 areas) feature selection (top: males, bottom: females) depicting significantly enhanced clustering efficiency. (C) A multinomial logistic regression model with Elastic net regularization was employed to predict categorical sickness states using only the Boruta-selected regions. (D) Normalized confusion matrices (left: males, right: females) showing the classification accuracy of the model. Clear diagonal dominance corresponds to correctly classified cases, demonstrating the model’s strong discriminatory power. (E) and (F) Mean Coefficient Profiles: Mean model coefficients for the 36 selected brain regions in males (E) and the 25 selected regions in females (F) for each sickness state. Coefficients were averaged across the 100 repeated cross-validation runs. A positive coefficient indicates that increased FOS activity in that region drives the prediction toward that specific sickness state, while a negative coefficient suggests that suppression of its activity increases the likelihood of the associated state. Bars depict ±SEM. Full list of brain areas in (E) and (F) included in Table S1 and Table S2, respectively.

Next, we employed a multinomial logistic regression model with Elastic Net regularization to predict categorical sickness states using only the brain regions selected by the Boruta feature selection algorithm (Fig. 4C). The Elastic Net penalty (combining L1 and L2 terms) was chosen specifically for its robustness in handling the high-dimensional, potentially correlated nature of FOS data (Fig. S4B). Next, we generated confusion matrices (Fig. 4D) and evaluated class-specific prediction probabilities for each sickness state (Fig. S4C&F). This analysis confirmed the model’s strong discriminatory power: high prediction probabilities were consistently assigned to test samples belonging to their true sickness state, and these probabilities dropped markedly for samples originating from different conditions (Fig. S4C&F). The confusion matrices further illustrate that the model accurately distinguished between sickness states, with clear diagonal dominance corresponding to correctly classified states (Fig. 4D). We also noted that the model exhibited higher confusion between the DSS and STAg sickness states compared to any other pair of conditions in both the male and female datasets. A higher confusion was also observed between LPS and pI:C in males. This specific pattern of misclassification mirrors our earlier observations across the behavioral, physiological, and peripheral cytokine analyses. Collectively, these results demonstrate that pathogen-specific sickness states evoke distinct patterns of response observable at both the organismal and cellular (FOS expression) levels.

We extracted and averaged the class-specific model coefficients across the 100 repeated stratified k-fold cross-validation runs. These coefficients not only rank each feature’s contribution but also convey directional information: a positive coefficient indicates that increased FOS activity drives the prediction toward that specific sickness state Fig. S4D&G), whereas a negative coefficient reflects an association with an alternative or baseline state (Fig. 4E&F, Table S2&3).This approach produced consistent coefficient profiles for each sickness state, enabling us to further reduce the number of functionally relevant brain areas for each sickness state for both males and females.

To further refine our interpretation of feature importance, we computed a hybrid importance score defined as abs(SHAP) × B, highlighting brain regions that are both impactful and consistent in their influence (Fig. S4E&H). This measure revealed class-specific patterns of directional prediction; for example, area postrema (AP) exhibited a strong positive effect on the PLA state (Fig. 4E&F, Fig. S4E), whereas superior colliculus (SCdw) contributes negatively to STAg classification, suggesting that suppression in SCdw is predictive of STAg (Fig. S4E).

This analysis demonstrates that each sickness state possesses a distinct and reliable neuronal marker. The convergence of high predictive accuracy (multinomial regression), clear clustering separation (t-SNE), and the identification of robust, class-specific neuronal activity patterns (SHAP) suggests a unique FOS signature for every sickness state. Finally, this and other analyses supported the conclusion that the effects of STAg and DSS, on one hand, and LPS and pI:C, on the other hand, form two separate clusters of sickness states, whereas PLA2 appears the most distinct from other models.

### Different immune challenges induce distinct molecular changes in the preoptic area of the hypothalamus

The identification of changes at the organismal and cellular scales substantiated the differential coding of immune challenges in the brain and prompted us to assess whether these differences were also reflected at the molecular level. We focused our experiments on the hypothalamic preoptic area, for which molecularly defined cell types associated with innate social behaviors and homeostatic functions have been delineated ^49,50^.

In an initial experiment, we performed bulk RNA sequencing of the hypothalamic preoptic area of mice treated with DSS or injected with LPS, pI:C, PLA2, or STAg. All conditions except injection of STAg showed a large number of distinct and overlapping differentially expressed genes compared to saline-injected controls, suggesting molecular changes that are unique to each sickness model. Importantly, we found that many of the differentially expressed genes corresponded to transcriptomic programs known to be involved in the response to different pathogens (Fig. S5A&B). For example, numerous interferon-stimulated genes, such as *Oasl1, Ifit1,* and *Isg15,* were upregulated in pI:Cinjected mice, and to a lesser extent in LPS-injected mice (Fig. S5B&C). This reflects the well-established role of interferon responses in antiviral immunity and activation of the non-canonical, toll-like-receptor 4 pathway downstream of LPS detection^22,51^. We also found several genes encoding antibacterial effectors to be upregulated in LPS-injected mice, such as *Il1b*, *Lcn2*, and *Ccl2* (Fig. S5D).

To gain a more granular understanding of sickness-induced gene expression in the POA, we used the imaging-based spatial transcriptomic platform, multiplexed error-robust *in situ* hybridization (MERFISH). We and others have used MERFISH to assess spatially-resolved single-cell gene expression in a number of contexts including social behavior, age-associated inflammation, and fever during sickness to identify molecularly distinct neuronal cell types and their transcriptomic changes across different biological contexts^12,50,52^. MERFISH probe libraries primarily were designed with the following goals (1) define the inflammatory landscape of the POA during homeostasis and sickness, (2) identify differentially expressed genes, with a particular emphasis on immune genes, across neurons and glia during sickness, and (3) define the neuronal cell-types that are most affected by different sickness models. In pursuing this third goal we aimed to establish a resource that could help inform future studies on sickness behavior, neurodegeneration, and other contexts of inflammation in the brain by comprehensively profiling the immune landscape of neurons and glia in the POA.

We generated a 940-gene MERFISH spatial transcriptomic atlas of the anterior mouse hypothalamus (from the preoptic to paraventricular areas) across these conditions (Fig. 5A, Table S4.). We employed a ‘dual-library’ design in which one library was designed for maximally resolved cell identification and the other for profiling most immune-related genes of interest. Based on pilot RNA-seq experiments (Fig. S5A-D), we further included a select number of differentially expressed genes in the immune profiling library. For a subset of genes, predominantly comprising neuropeptides (*Oxt, Avp*, *Gal*, *Sst, Penk*, *Tac1*, *Tac2*) we anticipated high expression that would lead to optical crowding, hence they were instead included as additional sequential FISH stains (Table S4).

**Figure 5.**
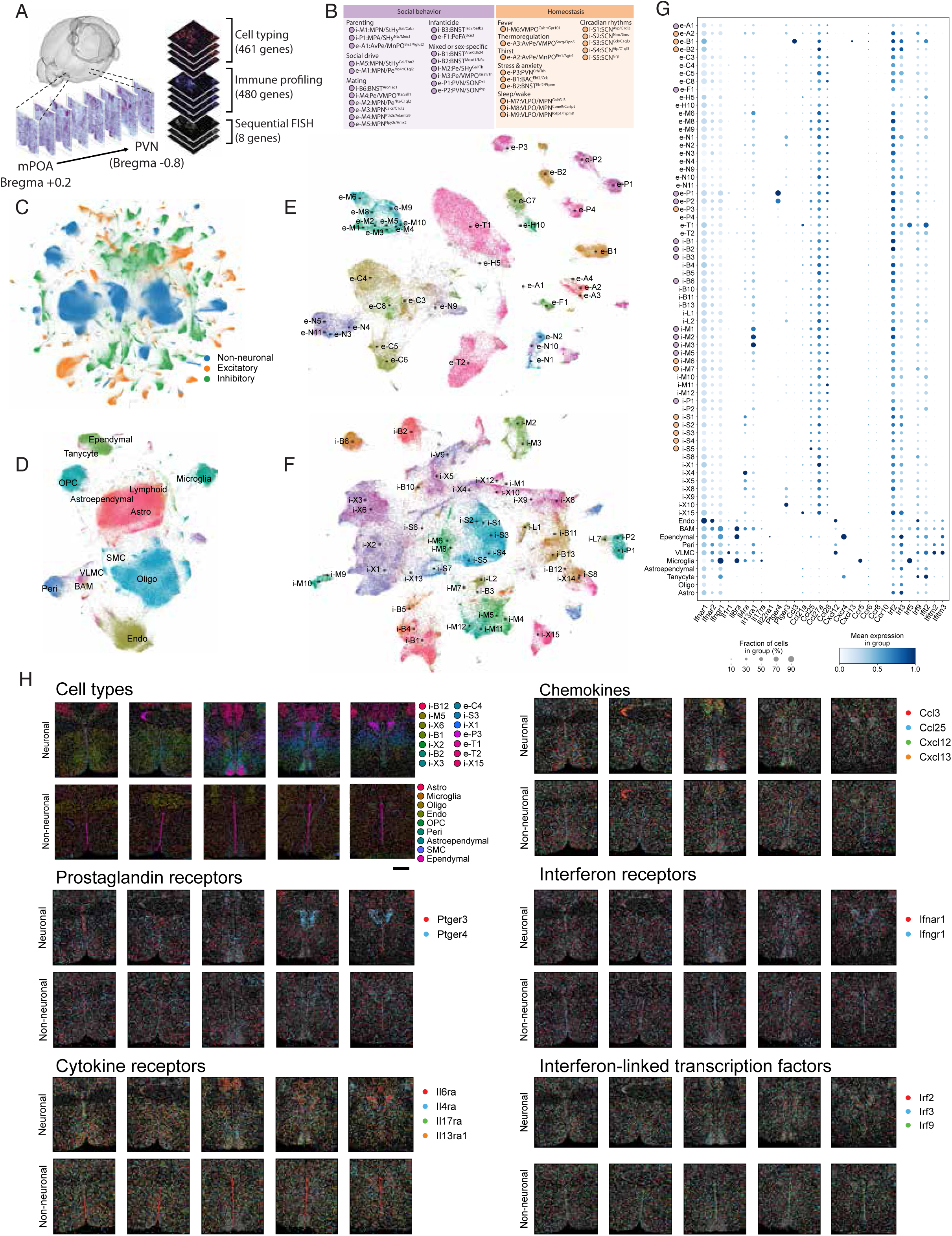
Defining the immune landscape of the preoptic and paraventricular hypothalamus. (A) Schematic of hypothalamic region of interest and MERFISH library design. (B) Functional annotations of cell types involved in regulation of physiology and social behavior (C-F) UMAPs of mapped cellular populations. (G) Expression of select immune genes (H) Spatial localization of cell-types and select immune genes across neurons and non-neuronal cells. For brevity, the legend shows cell types with greater than 1000 cells in the displayed sample. Scale = 500µm.

Across 16 animals and 173 hypothalamic sections we recovered 1.95 million high-quality cells, resolving a remarkable diversity of neuronal and non-neuronal cell types (Fig. 5BF). To provide a comprehensive descriptor for each cell, we used deep learning to integrate our MERFISH dataset with the Allen Brain Atlas and a P65 hypothalamus singlenucleus RNA-seq atlas^49,53^.

### Constitutive expression of immune genes in the POA and PVN

To establish a comprehensive molecular atlas of immune signals in the POA and PVN, we first mapped their cell type and spatial distributions at the normal health baseline. We began by performing label-transfer to assign cell-types identified in our dataset to celltype annotations previously defined by our lab according to their location and biological function (Fig. 5B-F, Fig. S5E-J)^49,50^. This allowed us to assess the transcriptome of numerous hypothalamic cell-types involved in the regulation of social behavior and homeostatic functions altered in sickness, such as social drive, body temperature, and sleep, as well as non-neuronal cells (Fig. 5B-F). We next assessed the expression of immune genes in individual cell-types in saline-injected mice to gain an understanding of the “inflammatory landscape” of the POA and PVN at baseline. By considering expression of genes above the blank identification threshold, we found numerous immune genes to be constitutively expressed across the POA and PVN in saline-injected mice in several intriguing functional categories (Table S5 & Fig. 5G). Genes involved in type-1 and type2 interferon signaling, such as *Ifnar1, Ifnar2, Ifngr1* and *Irf3*, were constitutively expressed across all cell types, and more highly in glia than in neurons. Other genes were more restricted to individual cell types: *Il13ra1*, involved in type-2 immunity, was preferentially expressed in i-M1, i-M2, and i-M3, MPN neurons that regulate social behavior, and the interferon pathway-related genes, *Irf5* and *Irf9*, were enriched in e-T1, excitatory neurons in the paraventricular thalamus (PVT) (Fig. 5G). *Ptger4*, which encodes the receptor for prostaglandin E2, was highly expressed almost exclusively in e-P1 and e-P2, neurons in the PVN that produce oxytocin and vasopressin, respectively (Fig. 5G). Non-neuronal cells also exhibited constitutive expression of numerous immune genes. Border-associated macrophages (BAMs), ependymal cells, and microglia showed enhanced *Ifngr1* expression compared to neurons, in addition to high expression of *Il6ra*, and to a lesser extent, *Il4ra* and *Il13ra1*, suggesting that these cell types have the capacity to respond to both type-1 and type-2 cytokines. Other genes, particularly chemokines and chemokine receptors, showed more cell type-specific expression patterns, rather than a shared pattern of expression across cell types, as observed in neurons for *Ccl25, Ccl27a,* and *Ccl28*. Among non-neuronal cells, we observed higher and more widespread expression of immune genes in BAMs, ependymal cells, pericytes, vascular leptomeningeal cells, and microglia (Fig. 5G).

We next assessed the expression of receptors for cytokines we found to be induced in serum across sickness states (Fig. 1B, Fig. S1C), as well as for prostaglandins, which are known to regulate a number of sickness behaviors.^15,16,54^. Prostaglandin receptor *Ptger3* showed widespread expression in the areas we assessed, while *Ptger4* showed strong spatial localization in the PVN (Fig. 5H). Among cytokine and interferon receptors found to be differentially upregulated in the serum across sickness states, *Il6ra, Il4ra, Il17ra,* and *Il13ra1,* as well as *Ifnar1* and *Ifngr1* were constitutively expressed. Certain receptors, including *Il6ra* and *Il4ra* showed spatial specificity. *Il6ra* was enriched in PVT and CRH-producing PVN neurons, e-P3, as well as in cells lining the third ventricle (Fig. 5G&H). *Il4ra* was enriched in the suprachiasmatic nucleus (SCN), particularly in i-S1 and i-S2, which are involved in circadian control (Fig. 5G&H). In addition to high expression of the type-1 interferon receptor *Ifnar1*, we saw widespread expression of transcription factors involved in interferon signaling, including *Irf2, Irf3,* and *Irf9* (Fig. 5H). Intriguingly, we observed highly specific expression of chemokines, such as *Cxcl12,* in the paraventricular thalamic (PVT) e-T2 neurons, and *Ccl3/Cxcl13* in the bed nucleus of the anterior commissure (BAC) e-B1 neurons. Together with previous studies from our lab and others showing that chemokines can directly signal to hypothalamic neurons and regulate their activity ^15,55,56^, this suggests a general role for chemokine signaling in the brain, potentially independent of its canonical role in chemotaxis. Together, these observations indicate that across many cell types involved in homeostasis and social behavior, both neurons and glia are poised to respond to diverse immune signals in broad as well as more spatially and cell type restricted patterns.

### Cell-type specific inducible gene expression

Next, we sought to understand the extent to which different cell types were affected by peripheral immune challenge. For this purpose, we defined the degree to which the transcriptome of individual cells was differentially regulated under the different sickness states explored above (Fig 6A). As expected, periphery-facing populations, such as the endothelial cells, were the most perturbed, with the highest number of DEGs. Of the neuronal populations, the PVN e-P1/2/3 neurons were most affected, likely reflecting the role of the PVN as a master regulatory region responding to molecular sickness cues, and further supported by our brain-wide analysis showing strong FOS induction in this region (Fig. 3D; Fig. 6A). In the POA, several populations of interest involved in regulating many sickness behaviors showed differential expression, including i-M6:VMPO^Calcr/Gpr101^, previously identified as regulating fever^15^, i-M5:MPN/StHY^Gal/Fbn2^, recently identified as regulating social homeostasis^42^, i-S1:SCN^Avp/C1ql13^, involved in circadian rhythm^49^, and eA2:AvPe/MnPO^Etv1/Agtr1^, which regulate thirst^57^ (Fig. 6A).

**Figure 6:**
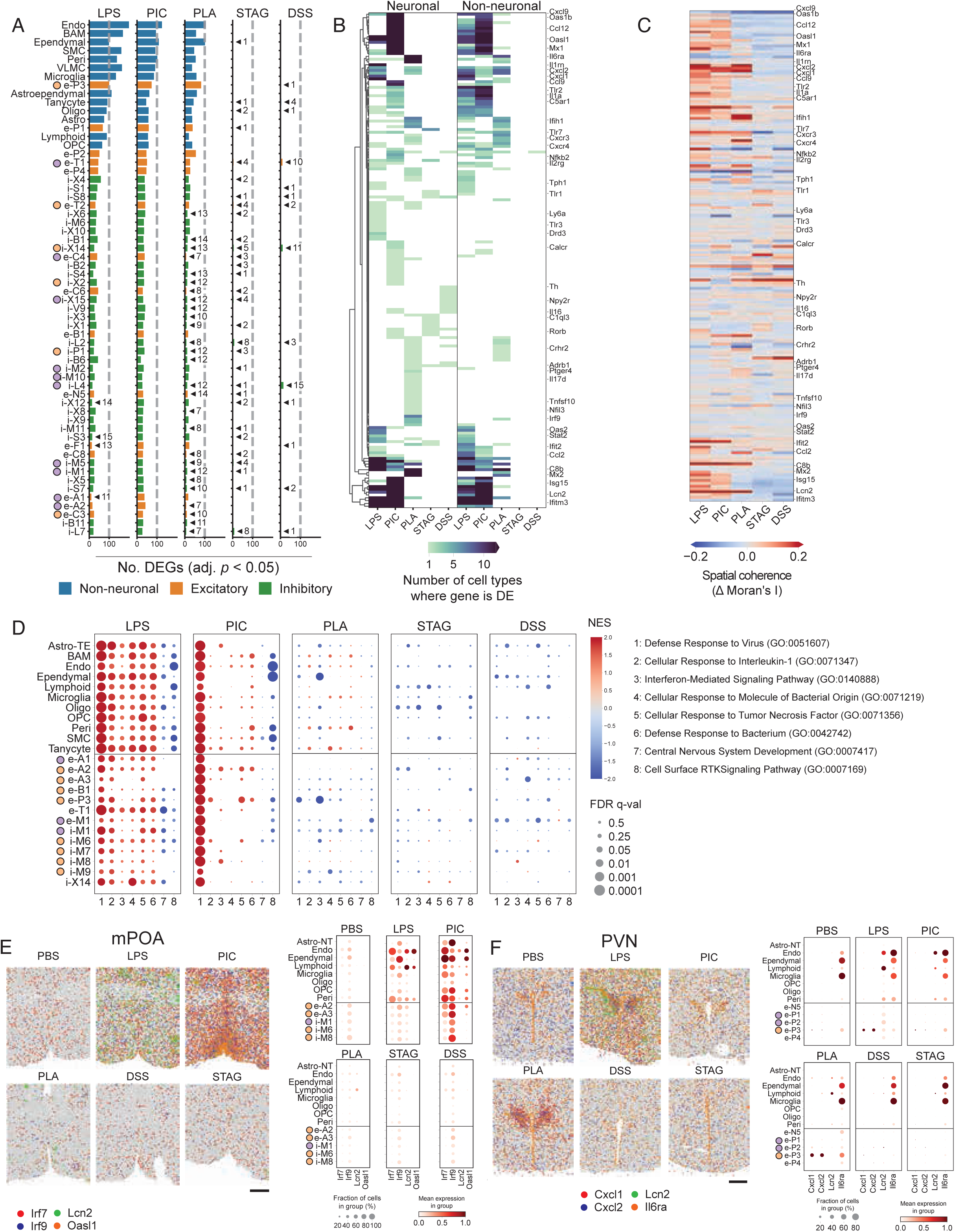
Differential responses across neurons and non-neuronal populations under different immune challenges. (A) Number of pseudobulk DEGs within a given cell type, with adjusted p < 0.05 and log2 fold change of 1 or greater. (B) Cell type specificity of differentially expressed genes, separated by neuronal and non-neuronal types. A more stringent threshold of adjusted p < 0.01 and a log2 fold change > 2 is used to mitigate the impact of spurious hits. (C) Spatial coherence of differentially expressed genes. The delta Moran’s I metric is measure the difference in spatial autocorrelation between treated and control animals. Higher values suggest focal or region-specific upregulation, while lower values suggest broad upregulation. (D) GSEA analysis showing a select subset of terms and cell types. (E) Upregulation of immune genes in the mPOA. (F) Upregulation of immune genes in the PVN. Scale = 500µm.

Across these populations, we further asked which DEGs had cell type-specific versus broad sickness-induced upregulation. As a simple heuristic, we counted the number of cell types in which a given gene was observed to be upregulated (Fig. 6B). Consistent with our bulk RNA-seq data, genes associated with antiviral immunity via type-1 interferon signaling were upregulated across many neuronal and non-neuronal populations in pI:C- and LPS-injected mice, such as *Oas1b, Oasl1, Oasl2, Mx1, Irf9, Mx2, Isg15, Irf7, Ifit1,* and *Ifit3* (Fig. 5B, Fig. S6A, Table S5). We also saw changes in chemokine expression, including *Cxcl9, Ccl12, Cxcl2,* and others, supporting the hypothesis of a previously unappreciated role for chemokine signaling in the brain (Fig. 6B&F, Fig. S6A, Table S6). To further identify genes that are upregulated in a region-specific manner, we employed a differential spatial autocorrelation analysis. Here, the Moran’s I statistic served as a proxy for regional specificity. Several genes showed regionalized gene expression changes, including several interferon-stimulated genes (ISGs) and chemokines (Fig. 6C, Table S6).

Next, we looked at functional categories enriched in cell-type-specific differentially expressed genes using gene set enrichment analysis (GSEA) and uncovered interesting patterns. As expected, pI:C-injected mice showed upregulation of genes involved in antiviral immunity across all cell types. LPS injection elicited broad gene expression changes across relevant functional categories, including antibacterial defense, IL-1 signaling, and TNF signaling (Fig. 6D). Non-neuronal cells showed a greater degree of gene expression changes across several gene ontology terms, while certain terms were more enriched in certain neuronal cell-types than others. For example, e-A2, involved in the regulation of thirst, showed enrichment of genes involved in antiviral immunity in both LPS- and pI:C-injected mice. i-X14, a cell type whose function has not been defined, showed enrichment of genes involved in antiviral immunity and antibacterial immunity in LPS-injected mice (Fig. 6). We also noted downregulation of genes related to CNS development in non-neuronal cells in LPS- and pI:C-injected animals.

Finally, we examined the spatial localization of cell-type-specific gene expression changes in cells involved in regulating processes that are altered during sickness. pI:Cinjected animals showed regionalized upregulation of the interferon-stimulated gene *Oasl1*, which was upregulated in endothelial and ependymal cells (Fig. 6E). *Lcn2*, involved in antibacterial immunity, was upregulated most strongly in LPS-injected animals, but predominantly expressed in non-neuronal cells, including pericytes, lymphocytes, and endothelial cells (Fig. 6E). We also observed intriguing spatial patterns of differential gene expression in the PVN. One of the most striking examples was the strong upregulation of *Cxcl1* and *Cxcl2* in cotricotropin-releasing hormone (CRH) neurons, in PLA2-injected animals and, to a lesser extent, in LPS-injected animals, perhaps associated with neutrophil infiltration (Fig. 6F). As expected, pI:C-injected animals showed strong upregulation of interferon-stimulated genes across numerous celltypes (Fig. S6).

Altogether, this analysis identified widespread expression of immune-related genes expressed in the POA and PVN, and showed how their expression differs across cell type, region, and sickness state. These data characterized pathogen-specific sickness states at the molecular level and represent a rich resource of comprehensive expression profiles of immune genes across cell types in this region.

## Discussion

We have uncovered distinct pathogen-specific sickness states triggered during different immune challenges and have characterized the corresponding changes at the organismal, cellular, and molecular levels (Fig. 7). Our findings indicate that, as in the peripheral immune system, the brain encodes distinct representations of different types of pathogens and elicits distinct behavioral repertoires. To date, a majority of studies on sickness behavior have relied on injection of mice with LPS to induce sickness, a robust model that has led to numerous and significant discoveries on the neural regulation of sickness behavior. However, the lack so far of thorough investigation of sickness in other infection models and inflammatory conditions, has limited our ability to comprehensively understand the diversity of neuroimmune interactions that can lead to sickness. In periphery, the generation of tailored immune responses is important for host defense – the elimination of an extracellular bacterial infection that produces enterotoxins requires different effectors than the elimination of a virus that has hijacked host cell translation machinery. This organization of immunity at the cellular and tissue scales enhances host fitness. A similar degree of specificity may exist at the organism scale, manifesting as different sickness states.

**Figure 7.**
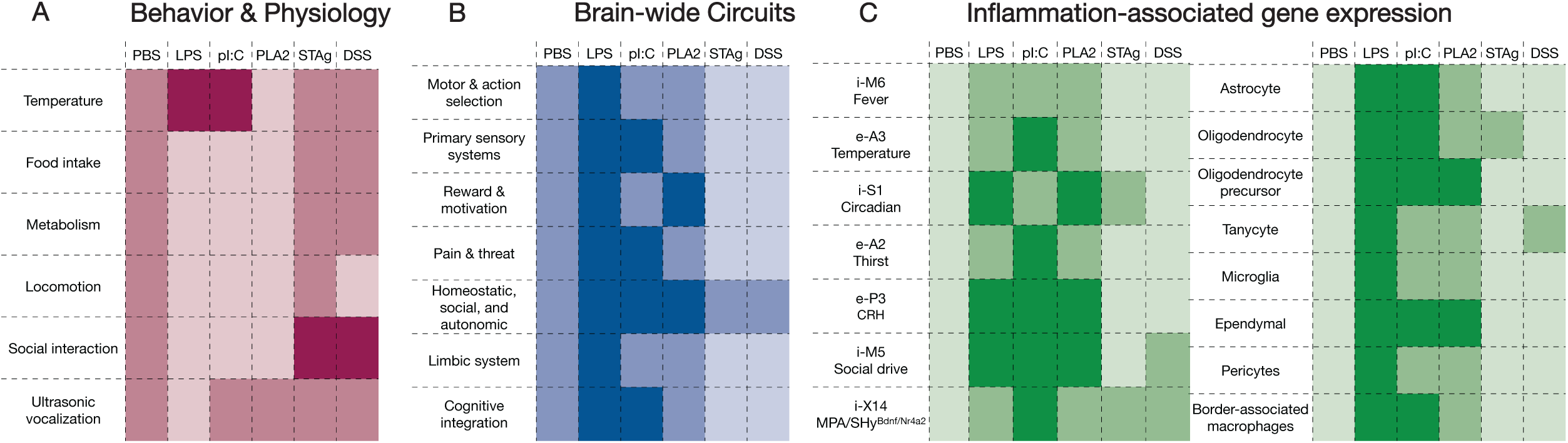
Pathogen-specific sickness states across biological scales. (A) Organismal responses to different sickness states assessed via physiological and behavioral screening. (B) Cellular responses to different sickness states assessed via whole-brain FOS staining. (C) Molecular responses to different sickness states in specific neuronal cell types (left panel) and glia (right panel) assessed via spatial transcriptomic analysis.

In this study, we used five different sterile immune challenges that model bacterial, viral, and parasite infection, as well as allergic immunity and colitis, to understand the generation of pathogen-specific sickness states in mice. We performed analyses at multiple biological scales – organismal, cellular, and molecular – and gained unique insights about how different sickness states are manifest at each level (Fig. 7). Across biological scales, we observed distinct and overlapping sickness states that form clusters based on phenotypic similarity: one including LPS and pI:C (modeling bacterial and viral infection), one including STAg and DSS (modeling parasite infection and colitis), and the third, PLA2 (modeling allergic inflammation), most distinct from the other conditions (Fig. 7).Though the observed similarity between STAg and DSS was not expected, other similarities are consistent with the distinct and overlapping peripheral immune responses to bacterial and viral pathogens and suggests that the distinction between type-1 and 2 immunity is conserved between the periphery and the brain. Indeed, the effect of PLA2 differed from other sickness states across behavioral, physiologic, cellular, and molecular analyses (Fig. 7). Surprisingly, responses to STAg and DSS differed significantly from other sickness models as well, particularly with respect to social behavior. In both immune challenges, mice exhibited increased prosocial behavior, pursuing interactions with their cagemates to a degree similar to saline-injected mice after social isolation (Fig. 7A). Our brain-wide FOS and spatial transcriptomic analyses revealed more modest responses to STAg and DSS than observed in other models, suggesting that these inflammatory models elicit milder states of sickness than others, despite their significant and unexpected impact on social behavior (Fig. 7B&C). Additionally, *Toxoplasma gondii* infection (modeled by STAg) and colitis (modeled by DSS) exhibit dynamic changes over longer time periods than assessed here, and future work focused specifically on these conditions could reveal temporal changes in the severity of sickness and its representation in the brain.

In addition to differences in sickness across immune challenges, our studies also revealed intriguing sex differences, most notably in pI:C- and PLA2-challenged mice. Female mice injected with PLA2, exhibited profound hypothermia, whereas males showed a much milder response. This observation is consistent with numerous reports of increased sensitivity to allergic or type-2 sensitization in females, for which hypothermia is a common readout in mouse models of allergy^44^. In comparison with males, PLA2injected females also displayed a minimal change in whole-body metabolism as assessed by respiratory exchange ratio. These results suggest interesting sex-dependent effects of acute type-2 immune challenge on metabolic and thermogenic systems.

Our in-depth single cell transcriptional study of the POA and PVN with MERFISH revealed unexpected features. Many immune genes were identified as constitutively expressed by distinct neuronal and non-neuronal cells, suggesting that specific cell populations, including neurons with known functions in homeostasis and social behavior control (Fig 7C) are poised to differentially respond to immune challenges. The specific repertoire of gene expression changes identified across conditions and by distinct neuronal and nonneuronal cell types further illustrates the complexity of the brain responses to immune challenges during sickness. Determining whether these changes represent adaptive or pathological responses provides an exciting future area of investigation. More generally, our results illustrate how sickness can differ both between sexes and across immune challenges. This establishes a rich landscape for future investigations into the causal relationships between cellular and molecular changes in the brain and sickness behavior across contexts.

In sum, our findings demonstrate that neuroimmune interactions generate pathogen-specific sickness states at the organismal, molecular, and cellular level, and challenge the notion of a generic representation of pathogens in the brain. This indicates that, like the immune system, the brain generates distinct responses to different kinds of immune challenges, resulting in distinct and overlapping changes in physiology and behavior. In the immune system, these context-dependent responses play an important role in generating the appropriate host defense strategy The parallels discovered here in the distinct responses of the immune system and brain to different classes of pathogen-specific triggers suggest that pathogen-specific sickness states may enhance host defenses. Understanding the role of the brain in host defense therefore requires updating our model of the brain response to infection as diversified across pathogens.

Our work also has important implications for understanding how chronic infection and neuroinflammation can contribute to disease. It is now well appreciated that inflammation plays a role in the pathogenesis of numerous noncommunicable diseases, including many that affect the brain. Our immune atlas of the POA and PVN, which is scalable to other brain areas, may provide clues into which specific inflammatory signals could contribute to altered brain function in different regions. The cell-type and region-specific expression of different inflammatory mediators and their receptors that we have reported here also suggest that an updated understanding of "neuroinflammation” as more specific than previously understood could help reveal underlying mechanisms of brain diseases. In addition to neurodegeneration and neurodevelopmental disorders, chronic infections are now well-understood to have profound effects on behavior and brain function. Post-acute inflection syndromes, of which long-COVID is an archetypal example, represent a major public health challenge and cause of disability for millions of patients. Though the mechanisms underlying these conditions remain poorly understood, the comprehensive behavioral, physiological, cellular, and molecular phenotyping we report here in pI:C challenged mice could help uncover how the antiviral immune response influences brain function.

Our work points to numerous future directions that will deepen our understanding of the neuroimmune interactions that underlie sickness behavior. Our molecular atlas of immune signals in POA- and PVN-treated mice highlights dozens of candidate immune mediators that may affect the function of defined cell types in these brain areas. Evaluating these candidates experimentally will likely uncover new molecular and cellular mechanisms that regulate sickness behavior during diverse immune challenges. Though we sought to be as comprehensive as possible in our selection of non-infectious immune challenges, our work here does not fully capture the landscape of inflammatory and infectious triggers that can influence physiology and behavior. For example, a recent study demonstrating the role of interleukin-6 in cancer cachexia^58^, a debilitating wasting syndrome, points to the importance of studying sickness-like behaviors in non-infectious inflammatory conditions such as cancer, metabolic disease, and aging. Finally, the proof-of-concept studies here leveraged robust, sterile models of infection and inflammation that can be used under biosafety level 1 conditions and therefore afford great flexibility in the application of downstream analyses and manipulations. However, in future work it will be important to evaluate pathogen-specific sickness states in *live* pathogen models, which better replicate the physiological conditions under which sickness is generated.

## Limitations of the study

The goal of the present work was to provide proof-of-concept evidence across multiple different experimental modalities that different sickness models elicit different phenotypes, and to establish a platform for understanding how host-pathogen interactions, across diverse pathogens, influence sickness behavior. However, all sickness models used in this study are sterile models of infection, not live pathogens, and therefore likely do not fully capture the complex temporal dynamics and host-pathogen interactions that would be observed in a true infection, nor do they enable us to address which changes we observe are adaptive versus pathological. It is also an inherent limitation of our approach that specific time points were selected for a given type of analysis as the responses to these perturbations are dynamic and thus would benefit from real time analysis such as activity imaging or *in vivo* electrophysiology to monitor activity changes in different sickness models. Furthermore, given inherent qualitative differences in the biomolecules and chemicals used to elicit sickness, we were unable to formally establish equivalent doses of different compounds. Inter-individual variability is also an intrinsic limitation of our models, and, indeed, has been leveraged by others to gain biological insight into protective physiologic mechanisms during infection^47,48^. The MERFISH studies used a curated library of genes of interest based on published literature that aimed to capture diverse cell-types and functionally relevant immune genes. Though we surveyed over 900 genes, this targeted approach is inherently biased, and it is possible that relevant genes were not captured in our library. Finally, our molecular experiments assessed gene expression, which may not faithfully represent protein levels across all genes analyzed in our investigations compared to proteomic approaches.

## Resource availability

### Lead contact

Further information and requests for resources should be directed to the lead contact, Catherine Dulac.

### Data and code availability

MERFISH data have been deposited with Data Dryad, currently available at: https://datadryad.org/share/LINK_NOT_FOR_PUBLICATION/yNoBDHMfZeG6y0LNDD bFQrGuDkeIHqOEdwH7_lF5BuU

Our pipelines for MERFISH processing and whole brain FOS prediction can be found at: https://github.com/Dulac-Lab/Brain-immune-interactions.

## Acknowledgements

Stacey Sullivan for assistance with animal husbandry and logistics. Drs. Allen, Osterhout, Giuliano, Zavatone-Veth, Jourjine, and Andreas for many helpful suggestions, advice, and assistance. We thank members of the Dulac lab and Dr. Richard Losick for helpful feedback on our manuscript. We thank LifeCanvas Technologies; the Beth Israel Deaconess Medical Center Physiology Core, including Emily Langmeyer and Drs. Banks and Berdan of the Harvard Chan Bioinformatics Core, Harvard T.H. Chan School of Public Health, Boston, MA, RRID:SCR_025373, for assistance with Bulk RNAseq analysis.

## Funding

This work was supported by: R01NS112399 and the Hock E. Tan and K. Lisa Yang Center for Autism Research at Harvard University to C.D.; R01GM143277 to J.R.M.; R01AI144369, R01AI158501, and Burroughs Wellcome Fund 1021330 to S.L.; Jane Coffin Childs Medical Research Awards grants 61-1749 to H.S.K; and a Hanna H. Gray Fellowship from the Howard Hughes Medical Institute to Z.A.S. Work by the Harvard Chan Bioinformatics Core was supported in part by the Harvard Catalyst: The Harvard Clinical Translational Science Center (National Center for Advancing Translational Sciences, National Institutes of Health Award UL 1TR002541). C.D. is an investigator of the Howard Hughes Medical Institute.

## Declaration of Interest

J.R.M. is an inventor of patents applied for by Harvard University and Boston Children’s Hospital related to MERFISH. J.R.M. is a co-founder and consultant of Vizgen, Inc. The other authors declare no conflict of interest.

## Supplementary Tables

**Supplementary Table 1. Activated brain areas during sickness. Related to Fig. 3 and Fig. S3.**

This table lists the acronyms and full names of brain areas shown in coronal section heatmaps in Figures 3 and S3.

**Supplementary Table 2. Top Boruta-selected brain regions contributing to classification in males. Related to Fig. 4.**

This table lists the 36 brain regions selected via Boruta feature selection (100 repetitions, top 5%) from the original 641 regions for male animals. These regions were used as input for downstream multinomial logistic regression with elastic net regularization. The table includes region names and final area identifiers after standardization. These features showed class-specific predictive value in distinguishing among sickness states (PBS, LPS, PIC, PLA, STAG, DSS).

**Supplementary Table 3. Top Boruta-selected brain regions contributing to classification in females. Related to Fig. 4.**

This table contains the 25 brain regions selected from the full feature space using Boruta feature selection (100 repetitions, top 5%) for female animals. These were the features used in the multinomial elastic net classifier to predict class membership across inflammation conditions.

**Supplementary Table 4: Sequences Associated with MERFISH. Related to Fig. 5**

The ‘MERFISH Probe Sequences’ contains a name and sequence for each of the template oligonucleotides used to create the MERFISH probes associated with combinatorial barcodes. The ‘Sequential FISH Sequences’ sheet contains the name and sequence of oligonucleotide probes used to target the gene and isoform id listed in the name for genes targeted with non-combinatorial, sequential staining. The ‘NBCTL1 Codebook’ and ‘NBIPL1 Codebook’ sheets contain the gene names, isoform ids, and binary barcode associated with each gene targeted by each of these barcode sets. Entries starting with ‘Blank’ represents barcodes not assigned to an RNA. The ‘Codebook-Readout Mapping’ sheet contains the bit number associated with each readout sequence for each library. The ‘Other Oligo Sequences’ sheet contains the name, sequence, and purpose for all other oligonucleotides used in MERFISH, library amplification, and sequential FISH.

**Supplementary Table 5: Gene Expression in Clusters Identified in MERFISH. Related to Fig. 5.**

The average log-normalized expression for all measured genes in each of the identified cell populations for mice treated with saline. Gene names are listed in the first row and cell type names are listed in the first column.

**Supplementary Table 6: Properties of Induced Gene Expression in Sickness States Measured with MERFISH. Related to Fig. 6**

The ‘Num DEG Cell Types’ sheet contains the number of cell types, split by Neuronal or Non-neuronal, in which each of the listed genes was differentially expressed. The ‘Delta Moran’s I’ sheet provides the measured differences between the Moran’s I averaged across all measured slices in each sickness state listed and that measured for the samples treated with PBS.

## Methods

### Animals

Mice were maintained on a 12 hour:12 hour dark-light cycle with access to food and water ad libitum. The temperature was maintained at 22 °C and the humidity was controlled at 35 +30/-10%. All experiments were performed in accordance with NIH guidelines and approved by the Harvard University Institutional Animal Care and Use Committee (IACUC). Young adult (7-9 weeks) C57BL/6J (JAX 000664) and FVB/NJ mice (JAX 001800) were used for were used for social behavior, locomotion, temperature, and food intake experiments. C57BL/6J mice were used for cytokine, FOS, RNA sequencing, and MERFISH experiments. Sample sizes were chosen on the basis of similar experiments from relevant studies.

### Sickness Models

#### DSS

Dextran sodium sulfate (MP Biomedicals; Lot#: S8364) is mixed with distilled water to achieve a 3% concentration and added to a water bottle^39^. Dosing was optimized based on the lot and animal facility to ensure effective induction of colitis symptoms over the course of 3-7 days. Body weight was measured daily over the course of administration. Control mice also received a water bottle with similar volume without DSS to control for the change of water access from rack spout to water bottle in the cage hopper.

#### PLA2, pIC, LPS

Mice were injected intraperitoneally with 100 ug of Bee venom phospholipase A2 (SigmaAldrich), 50 ug/kg of lipopolysaccharide (Sigma Aldrich), and 10 mg/kg of Polyinosinic-polycytidylic acid (Invivogen) 4 hours prior to each experiment^15,59^. Doses were optimized to elicit a robust immune and behavioral response while still enabling mice to locomote. Based on time course cytokine levels in each model, an incubation period of four hours was chosen.

#### STAg

*Toxoplasma gondii* (Pru strain) was first grown in fibroblasts in cell culture. The host cells were lysed and filtered, yielding a solution containing only parasites. Parasites were then killed and lysed by five subsequent rounds of snap-freezing in liquid nitrogen followed by rapid thawing at 37 °C. After the freeze–thaw cycles, the mixture was sonicated to further lyse the parasites. The samples were then centrifuged to pellet the insoluble debris, and STAg was collected from the soluble supernatant.^35^ Mice were injected intraperitoneally with 25µg of STAg for each experiment.

### Behavior Assays

All behavioral experiments were performed during the dark cycle in a room illuminated by infrared or red light. Mice were habituated in the room for 10-20 minutes prior to beginning experiments. All behaviors occurring during the behavior period were recorded using a multi-camera surveillance system (GeoVision GV-VMS software and GV-BX4700-3V cameras).

#### Social Isolation / Reunion Assay

Female littermate mice were group housed (2-4 mice) for at least one week prior to isolation. Females were used due to their sensitivity to isolation and to mitigate aggressive behavior that often occurs during interactions between male mice^42^. Before isolation, two socially housed cage mates were placed in a new cage with fresh bedding and recorded for 10 minutes to measure the baseline social interaction (ad lib). One mouse was then isolated in a new cage for 4 days while the other mouse was kept socially housed. Following 4 days of isolation, the experimental mouse and the socially housed cage mate are returned to a clean cage with fresh bedding and recorded for 10 minutes to measure the social rebound.

#### Behavioral Analysis

Behavior videos were manually scored using the Observer XT 11 software (Noldus Information Software) to identify predetermined social behavior actions. Only behaviors initiated by the experimental mouse were analyzed. Social interactions were characterized by the following, mutually exclusive behaviors: sniffing, approaching/chasing, head-to-head contact, crawling (underneath the other mouse), allogrooming, and mounting. Mice were characterized as non-social when they were not initiating these behaviors.

#### Locomotion

Mice were injected intraperitoneally with saline, LPS, PLA2, STAg, or pIC, except for a cohort which received DSS in a water bottle 3 days prior to experiment. After 1 h of habituation, mice were placed in a new cage and recorded for 20 min. Tests were recorded using a Geovision surveillance system, and the total distance travelled, average velocity, and center point tracking and heatmaps were measured and generated using Ethovision XT 13.0 (Noldus Information Technology). To improve the reliability of automated tracking, a video segmentation model (specifically SAM2; https://arxiv.org/abs/2408.00714) was used to increase contrast between each mouse and the background in videos of behavior. In the first video frame containing a mouse, researchers annotated 3-5 points inside the image of the mouse, from which a segmentation mask was derived and propagated to later video frames. Finally, each segmentation mask was recolored.

#### Food Intake

All experimental mice were deprived of food for 22 hours before the experiment. After 1 h of habituation, mice were injected intraperitoneally with saline, LPS, PLA2, STAg, or pIC, except for a cohort which received DSS in a water bottle 3 days prior to experiment. A pre-weighed amount of regular chow was returned to the cage at the time of injection.

The weight of remaining chow was measured 4 h later.

#### Temperature

For intermittent temperature recordings, implantable temperature transponders (TP-1000 Temperature Programmable transponder from Avidity Science) were used to approximate basal body temperature. Mice were anaesthetized using 2.5-3% isoflurane. Transponders were inserted under the skin below the neckline in the upper abdominal region to the right of the midline. Hair was shaved and skin was disinfected before transponder insertion. Mice recovered for 1 week before experiments. Body temperature was measured immediately before behavior experiments using a handheld transponder reader (DAS-8037 Wireless Reader System, Avidity Science).

#### Metabolic cages

For metabolic cage experiments, animals were individually housed in Promethion High-Definition Multiplexed Respirometry Cages (Sable Systems International). After 48 hours of acclimatization, mice were injected with indicated compounds, or exposed to DSS in the drinking water, and monitored for VO2 and VCO2 an additional 24 hour (acute models) or 72 hours (DSS).

### Ultrasonic vocalizations

#### USV recording

Female FVB/NJ mice were exposed to the reunion assay. Following four days of isolation, the experimental mouse was injected intraperitoneally with saline, LPS, PLA2, STAg, or pI:C, unless it was part of a cohort given DSS to drink over the isolation period. The recordings were performed four hours later under red light. The experimental mouse and its socially-housed cagemate were placed in a new cage with fresh bedding and recorded for 10 minutes using an ultrasonic microphone (Ultrasound Gate CM16/CMPA; Avisoft Bioacoustics) placed 30 cm above the cage floor. The signal was converted to a digital format using an analogue-to-digital converter (Ultrasound Gate 116, Avisoft Bioacoustics) sampling at 500 kHz.

#### USV detection

USVs were detected with a temporal convolutional neural network (TCN), implemented by the software package Deep Audio Segmenter (DAS) v.0.32.5^60^. The model used was DAS’ tcn_stft model. It combines a TCN and a short time Fourier transform, which improves classification on rodent vocalizations according to Seinfarth et al.. All parameters used for training were default: np_epoch: 400, kernel_size: 16, nb_filters: 32, nb_hist: 1024, nb_pre_conv: 4, nb_conv: 3, learning_rate: 0.0001, batch_size: 32. The data used as input consisted of raw audio chunks of 1,024 samples, corresponding to approximately 2.048 ms at a 500 kHz sampling rate. This is substantially shorter than typical rodent USV but enables high temporal precision. The model was evaluated on its ability to label chunks as ’USV’ or ’noise’ (non-vocal segments). The model was trained using prediction-assisted labelling, which is an iterative process to speed up labelling. The first iteration was trained on a set of 350 manually annotated vocalizations, and four rounds of prediction-assisted labelling were performed. The final model was trained on 40,974 annotated vocalizations. Each labeled wav file was cut following a 60%/20%/20% train/test/validation split.

Vocalizations with low amplitude and low quality were still inconsistently detected by the model, and thus were manually annotated following visual inspection of audio recordings. Vocalizations shorter than 5 ms were excluded, resulting in a total of 83,608 individual ultrasonic vocalizations.

#### USV analysis

All analyses were performed in Python 3.12.3 and R 4.4.1. Frequencies lower than 20 kHz or higher than 125 kHz were omitted.

#### Feature extraction

Acoustic features and sound pressure levels were extracted using the spectro_analysis and sound_pressure_level functions from the R package warbleR (v.1.1.3)^61^ with the following parameters: ovlp: 25, mar: 0.005, wl: 1,024, harmonicity: F, Fast: T. The list of features computed can be found on http://marce10.github.io/warbleR/reference/spectro_analysis.html. In addition, four temporal metrics were calculated for each call n: onset_(n+1)-onset_n, onset_(n+2)onset_n, onset_(n+3)-onset_n and (duration_n)/(onset_n ).

#### Statistics

Python packages scipy (v.1.15.1) and scikit_posthocs (v.0.11.3) were used for statistical data analysis. Pairwise comparisons were performed using two-tailed Student’s t test for independent samples for parametric features and Wilcoxon rank-sum test for nonparametric features. Bonferroni correction was applied before plotting test results.

#### Prediction on features

Python package scikit-learn (v.1.6.1) was used to train a random forest classifier on all the computed features, with the following hyperparameters: n_estimators: 500, criterion: entropy, max_depth: 20, min_samples_split: 2, min_samples_leaf: 1, max_features: None. To avoid biasing the model towards over-represented classes, 50 forests were fitted to random balanced sub-samplings of the data, either 2,500 or 7,801 (the latter corresponding to every USV from the condition with fewest USVs). Smaller subsamplings affected model performance. Each sub-sampling was further divided into training (80%) and testing (20%) datasets, and a matching shuffled dataset was created by randomly shuffling all sub-sampled labels.

The metric used to measure predictive power was the F1-score, defined using precision=truepositive/(truepositive+falsepositive) and recall=truepositive/(truepositive+falsenegative) so that the F1 score corresponds to: F1score=2/(1/precision+1/recall). The F1 scores and feature importance are presented as mean ± SD. The confusion matrices are the averaged normalised confusion matrices over all iterations, meaning that within each cell is the mean proportion of "true label" predicted as "predicted label".

#### Spectrogram creation

Spectrograms were created from the raw audio signal of individual vocalizations, using a short-time Fourier transform with a segment length of 1,024 and overlap length of 512.

To focus the analysis on image features rather than duration differences, all spectrograms were linearly interpolated to the length of the longest articulate vocalization using the RegularGridInterpolator function of the Python package scipy. These spectrograms were finally sampled at 128x128 evenly spaced time and frequency points between onset and offset and between 20 kHz and 125 kHz, respectively.

#### UMAP embedding of spectrograms

The spectrograms were flattened, z-scored across samples, and embedded in two dimensions using uniform manifold approximation and projection with the Python package umap-learn (v.0.5.7) with default parameters^62^.

### In-Situ Hybridization (FISH)

Double-label fluorescence in-situ hybridization (FISH) was performed using a RNAScope assay V2 kit (Advanced Cell Diagnostics (ACD) according to the manufacturer’s instructions with a few exceptions. Freshly frozen brains were sectioned using a cryostat microtome at 14 μm and stored at −80 °C. The Allen Brain Atlas was used to determine locations for sectioning of the preoptic area based on anatomical landmarks. Slides were thawed and fixed in 4% PFA for 15 min followed by dehydration in 50%, 75% then 100% ethanol. Cells were permeabilized for 25 min at room temperature using Protease IV provided by ACD. Sections were then processed as suggested by the ACD protocol. Slides were imaged on a Zeiss Axioscan Z.7 at 10x using ZEN Blue 3.11 software (Zeiss) at the Harvard Center for Biological Imaging (RRID:SCR_018673). All probes were made by ACD. Quantification of marker expression was done using QuPath 0.3.2 that identified cells using the DAPI signal and identified cells with fluorescent signals corresponding to cFos, Trhr, and Mc4r respectively.

### Enzyme-Linked Immunosorbent Assay (ELISA)

Whole blood was harvested from mice by retro-orbital bleeding and plasma was isolated using lithium heparin coated plasma separator tubes (BD). Plasma samples were diluted 1:1. Multiplex cytokine analysis was performed by Eve Technologies Corporation (Calgary, Alberta, Canada) using the Luminex® 200™ system (Luminex Corporation/DiaSorin, Saluggia, Italy) with Bio-Plex Manager™ software (Bio-Rad Laboratories Inc., Hercules, California, United USA). The 44-plex analysis of mouse cytokines, chemokines and growth factors were measure using two separate Eve Technologies’ panels (Mouse Cytokine/Chemokine 32-Plex Discovery Assay® Array (MD32) (MILLIPLEX® Mouse Cytokine/Chemokine Magnetic Bead Panel Cat. #MCYTOMAG-70K, MilliporeSigma, Burlington, Massachusetts, USA); Mouse Cytokine/Chemokine 12-Plex Discovery Assay® Array (MD12) (MILLIPLEX® Mouse Cytokine/Chemokine Magnetic Bead Panel II Cat. # MECY2MAG-73K, MilliporeSigma, Burlington, Massachusetts, USA)). Assay sensitivities of these markers range from 0.3 – 30.6 pg/mL.

Samples with analyte concentrations outside of the range standard curve were assigned a value equivalent to either the highest or lowest concentration in the standard curve for that analyte. These were considered above or below the limit of detection as shown in each figure. For MIP-1b (CCL4), MCP-1 (CCL2), G-CSF, TARC (CCL17), and IL-10, values were too far outside the standard curve, so we manually applied a simple linear regression to approximate the observed values based on fluorescence.

### Whole brain FOS analysis

#### Tissue Preservation and Clearing, Immunolabeling and Imaging

Paraformaldehyde-fixed samples were preserved with SHIELD reagents (LifeCanvas Technologies) using the manufacturer’s instructions (Park et al., 2018). Samples were delipidated using LifeCanvas Technologies Clear+ delipidation reagents. Following delipidation samples were blocked using Antibody Blocking Solution (LifeCanvas Technologies) with 5% normal donkey serum (Jackson ImmunoResearch Laboratories Inc.) Samples were then labeled using the Radiant Buffer System (LifeCanvas Technologies) for primary labeling, followed by secondary labeling using SmartBatch+ Secondary Buffer system (LifeCanvas Technologies), both with eFLASH (Yun et al., 2019) technology which integrates stochastic electrotransport (Kim et al., 2015), using a SmartBatch+ device (LifeCanvas Technologies). After immunolabeling, samples were incubated in 50% EasyIndex (RI = 1.52, LifeCanvas Technologies) overnight at 37°C followed by 1 d incubation in 100% EasyIndex for refractive index matching. After index matching the samples were imaged using a SmartSPIM axially-swept light sheet microscope using a 3.6x objective (0.2 NA) (LifeCanvas Technologies).

#### Atlas Registration (Nuclear Dye)

Samples were registered to the Allen Brain Atlas (Allen Institute: https://portal.brainmap.org/) using an automated process (alignment performed by LifeCanvas Technologies). A nuclear dye channel for each brain was registered to an average nuclear dye atlas (generated by LCT using previously-registered samples). Registration was performed using successive rigid, affine, and b-spline warping algorithms (SimpleElastix: https://simpleelastix.github.io/).

#### Cell Detection

Automated cell detection was performed by LifeCanvas Technologies using a custom convolutional neural network created with the Tensorflow python package (Google). The cell detection was performed by two networks in sequence. First, a fully-convolutional detection network (https://arxiv.org/abs/1605.06211v1) based on a U-Net architecture (https://arxiv.org/abs/1505.04597v1) was used to find possible positive locations.

Second, a convolutional network using a ResNet architecture (https://arxiv.org/abs/1512.03385v1) was used to classify each location as positive or negative. Using the previously-calculated Atlas Registration, each cell location was projected onto the Allen Brain Atlas in order to count the number of cells for each atlas-defined region.

#### Data Preprocessing and Feature Selection

FOS intensity data from 641 discrete brain areas were first transformed using the YeoJohnson transformation to stabilize variance and approximate a Gaussian distribution. For feature reduction, we employed the Boruta feature selection algorithm, which uses a random forest classifier to compare real features against permuted "shadow" features^63^. This process was repeated 100 times, and the top 5% of features were retained, yielding statistically consistent subsets (e.g., 36 in males, 25 in females). This pre-selection step served to reduce noise and multicollinearity while preserving relevant biological variance.

#### Multinomial Logistic Regression with Elastic Net Regularization

To model sickness-state classification based on selected brain regions, we implemented a custom multinomial logistic regression model with elastic net regularization. The elastic net penalty, a linear combination of L1 and L2 norms, was chosen for its robustness in handling high-dimensional and correlated predictors. Model training included a nested cross-validation procedure: a 75–25% outer train–test split and an inner 4-fold crossvalidation for hyperparameter tuning (λ, α), repeated 100 times for stability.

#### Model Evaluation and Stability Analysis

Performance was assessed using metrics including accuracy and per-class mean squared error (MSE). To ensure robustness and interpretability, we computed model coefficients (B) and SHAP-style feature contributions across all iterations. Class-specific coefficients were averaged over folds, yielding stable effect size estimates with directional interpretation. Positive coefficients indicated that higher FOS activity in each brain region promoted classification into a particular sickness state, while negative coefficients implied suppression.

#### SHAP approximation and feature attribution

To interpret feature-level contributions, we used a linear SHAP (SHapley Additive exPlanations) approximation computed as:

SHAP(i,j,c) = X_std(i,j) × β(j,c)

This yielded signed SHAP values per feature, sample, and class, which were averaged across samples and folds to obtain stable estimates.

#### Visualization and Feature Interpretation

To interpret the model, we profiled the class-specific model coefficients averaged over 100 repetitions. This established stable coefficient profiles (Fig. 4E and F) that further refined the set of functionally relevant areas and revealed the directional influence of each region (positive β drives prediction toward a state; negative β suggests suppression is associated with that state).

For a fine-grained, instance-specific understanding, we augmented this analysis using a SHAP-style method. We prioritized features based on a hybrid metric, absolute SHAP value magnitude weighted by the stability of the regional coefficient β. This ensures that regions are selected not only for their per-sample influence (SHAP) but also for the robustness of their predictive power across repeated model fits. A high value in this combined metric, regardless of sign, indicates a strong and stable predictor.

### Bulk RNA sequencing & analysis

#### RNA isolation

Brains were dissected, frozen in optimal cutting temperature medium and placed at −80 °C. The POA and PVN surrounding regions were dissected on a cryostat at −15 °C using the Paxinos Adult Brain Atlas to determine locations for dissection based on landmarks. Tissue was collected from the anterior-most portion of MnPO to the posteriormost portion of the PVN. To achieve this, after slicing away anterior material, thin (10– 20 µm) sections were collected and examined under a dissecting microscope to identify landmarks. To collect tissue, we first used a razor blade to remove large blocks of tissue lateral to the lateral ventricles, and then sliced 150-µm-thick sections to collect roughly the ventral-most one-third of the brain into a 2-ml tube. Dissected tissue was placed in a 2-ml tube and kept at −80 °C until the day of RNA isolation. RNA was isolated using a Qiagen RNEasy Miniprep kit with on column DNAse digestion. Frozen tissue was homogenized in RLT buffer from Qiagen using a Kimble Disposable Pellet Pestle Tissue Grinder (Fisher K749515-0000). RNA isolation was performed according to manufacturer’s instructions.

#### Library prep & sequencing

RNA was measured using an RNA tape on the TapeStation 4200 (Agilent Technologies) to determine integrity. RNA concentration was measured using the ribogreen assay (Thermo Fisher Scientific) and all samples were normalized to 100ng. Polyadenylated mRNAs were captured using oligo-dT-conjugated magnetic beads (Watchmaker Genomics). Poly-adenylated mRNA samples were immediately fragmented to 300-400bp and stranded libraries were made by adding dUTP into the second strand with the Watchmaker mRNA library prep kit (Watchmaker Genomics) and X-Gen stubby adapters (Integrated DNA Technologies). Libraries were enriched and indexed using 11 cycles of amplification with the Equinox Master Mix (Watchmaker Genomics) and PCR primers which included dual 8bp index sequences to allow for multiplexing (Integrated DNA Technologies). All library preparation volumes were miniaturized to 1/5 reaction volume using the Mantis liquid handler (Formulatrix). Excess PCR reagents were removed through magnetic bead-based cleanup using Aline PCRClean Dx paramagnetic beads on a SciClone G3 NGSx workstation (Perkin Elmer). The resulting libraries were assessed using a 4200 TapeStation (Agilent Technologies) and quantified by QPCR (Roche Sequencing). Libraries were pooled and sequenced on one lane of a NovaSeq X Plus 25B flowcell using paired-end, 150 bp reads (Illumina, Inc.).

#### Differential expression analysis

Samples were processed through the nf-core rnaseq pipeline version 3.14.0^64^. Briefly, quality control of raw reads was performed using FastQC version 0.12.1^65^ and adapter and quality trimming was performed using cutadapt version 3.4 and Trim Galore version 0.6.7^66^. Trimmed reads were aligned to mouse genome GRCm38 using STAR version 2.7.9a^67^ and quantified using Salmon version 1.10.1^68^. Alignments were checked for evenness of coverage, rRNA content, genomic context of alignments (for example, alignments in known transcripts and introns), complexity, and other quality checks using a combination of FastQC, Qualimap version 2.3^69^, Samtools version 1.17^70^, MultiQC version 1.19^71^, and custom tools.

Differential expression by condition was called at the gene level using DESeq2 version 1.38.3^72^ using the counts per gene estimated by Salmon. Functional analysis of differentially expressed genes was examined for Hallmark pathways using the ClusterProfiler R package v4.6.2^73^. Plots were generated using the R packages ggplot2 v3.5.1, pheatmap v1.0.12, enrichplot version 1.18.4, EnhancedVolcano v1.16.0, and ComplexHeatmap v2.14.0. All of these analyses were performed using R version 4.2.3.

### MERFISH

#### Library design

MERFISH encoding probes were designed for 940 genes as previously described^50,74^. 30-nt target regions were designed, concatenated with 20-nt readout sequences comprising the assigned barcodes, and up to 72 such encoding probes were designed per gene, all using a previously described pipeline^50,74^. Genes were divided into two separate libraries comprising 463 and 481 genes. For each library, each gene was randomly assigned a binary barcode from a 24- or 26-bit Modified Hamming Distance (MHD4) code, respectively, where each gene is defined by four ‘on’ bits. For each target sequence, three of the four 20-nt readout sequences corresponding to the gene barcode were appended, alongside a T7 promoter and PCR amplification handles. Sequence libraries were ordered as a pool of oligos from Twist Biosciences.

Sequential single molecule (sm)FISH encoding probes for 8 genes (Oxt, Fos, Sst, Penk, Avp, Gal, Tac1, Tac2) were designed similar to above, except only a single readout sequence was concatenated to the target sequence. Sequential smFISH probes were directly ordered as an oPool from IDT. All encoding and readout probe sequences, as well as primer sequences for library amplification, are provided in **Supplemental Table 3**.

#### MERFISH Library amplification

Library amplification was performed as described previously^50,75^. Briefly, Twist Bioscience oligopools were resuspended and diluted 1:100 in IDTE pH 8.0 buffer (IDT #11-05-01) and amplified by qPCR (1:10 library, 5 µM each of the forward and reverse primers, 1× EvaGreen, 1× Phusion Hot Start Flex DNA Polymerase [NEB #M0535S]), stopping at a cycle during the exponential portion of the amplification process (typically <18 cycles). PCR products were purified on multiple Zymo III columns on a vacuum manifold as per manufacturer instructions. *In vitro* transcription (IVT) was carried out using a HiScribe T7 High Yield RNA Synthesis kit (NEB #E2040) supplemented with 1:30 Murine RNase inhibitor (NEB #M0314S) and an additional 100 µM of CTP and GTP. The IVT was incubated at 37 °C overnight. The resultant RNA was purified using magnetic beads.

Library ssDNA sequences were prepared by reverse transcription (120 µg IVT RNA, 1× RT buffer, 130 µM RT primer, 3.3 mM dNTPs, 1.5 µL Maxima H Minus Reverse Transcriptase [Thermo #EP0753], 1 µL RNAsin Plus [Promega N2615]) at 50 °C for at least 4 hours. Residual RNA was removed via alkaline hydrolysis. Specifically, 25 µL 0.5 M EDTA and 25 µL 1 M NaOH were added per 50 µL reaction, and tubes were incubated in a water-filled heat block at 95 °C for 15-20 minutes. Residual NaOH was quenched by addition of 30 µL 1 N HCl per reaction. The resultant ssDNA was purified on magnetic beads. Each preparation was quality controlled using Nanodrop measurements and denaturing PAGE.

#### Coverslip treatment

40mm-diameter #1.5 coverslips (Bioptechs #40-1313-03193) were cleaned and silanized as previously described^76^. Briefly, coverslips were immersed in a 1:1 mixture of 37% HCl and methanol for 30 minutes, washed with 70% ethanol, and allowed to dry completely at 60 °C. Cleaned coverslips were then silanized by immersion in chloroform containing 0.1% v/v triethylamine and 0.2% v/v allyltrichlorosilane for 30 minutes. The silane layer was allowed to fully cure by baking at 60 °C for at least 2 hours, followed by storage in a desiccated environment at least overnight. To support tissue adherence, coverslips were coated for at least 30 minutes with 1 mL poly-D-lysine (Thermo #A3890401) containing a 1:200 dilution of orange carboxylate-coated fiducial beads (Thermo #F8800). Excess solution was aspirated and coverslips were allowed to dry. Then, beads were affixed using 4% paraformaldehyde (PFA; Electron Miscroscopy Sciences #15714) in 1× PBS for 5 minutes. Coverslips were washed twice with water, then once with 70% ethanol and allowed to dry completely before use.

#### Sample preparation for MERFISH

Samples were prepared for MERFISH largely as described previously^50^. Fresh frozen mouse brains were cut to obtain roughly 3x3 mm hypothalamic tissue slices. In a typical sample, 14 µm-thick sections were collected from approximately Bregma +0.2 to -0.8 with an average spacing of 100 µm, spanning the major preoptic region to the paraventricular nuclei. Samples were immediately fixed with 4% PFA in 1× PBS for 20 minutes at room temperature. Subsequently, samples were washed thoroughly with 1× PBS to remove residual PFA, then permeabilized in 70% ethanol at 4 °C overnight. Samples were stored indefinitely at 4 °C until further use.

Samples were washed twice with 2× saline-sodium citrate (SSC) to remove residual ethanol, then equilibrated with 30% (v/v) formamide in 2X SSC. Coverslips were transferred to a humidified chamber system, and excess 30% formamide was aspirated. Tissue slices were covered with a 50 µL droplet of hybridization solution (30% formamide, 10% w/v dextran sulfate, 1 mg/mL yeast tRNA (Thermo #15401029), 8-10 µM each MERFISH encoding probe library (∼0.3 nM/oligo), sequential stain probe library (5 nM/oligo), 1 µM poly(A)-acrydite anchor probe, and 1% RNasin Plus (Promega #N2611) in 2× SSC). Pre-made hybridization mixes were stored at -20°C for several months with no observable decline in MERFISH quality. To improve hybridization quality, samples were heated at 60°C for 1 hour, before incubation at 37°C for at least 60 hours^77^. Nonspecifically bound probes were removed by washing twice with 30% formamide in 2× SSC at 47°C for 20-30 minutes each, then thrice briefly with 2× SSC at room temperature.

Samples were embedded for tissue clearing in a polyacrylamide hydrogel as described previously^76^. A gelation solution (4% v/v 20:1 acrylamide:bisacrylamide, 0.15% v/v TEMED, 0.3% w/v ammonium persulfate, 300 mM NaCl in 50 mM Tris-HCl pH 8.0) was prepared and briefly degassed, then samples were equilibrated with 2-4 mL gelation solution for 3 minutes. Subsequently, coverslips were inverted onto a Gel Slick (Lonza Bioscience #50640)-coated glass surface containing a droplet of gelation solution. Excess solution was removed, and gels were allowed to polymerize for 2 hours. Embedded samples were transferred to a new dish with 2 mL tissue clearing solution (2% SDS, 0.25% Triton X-100, 1% Proteinase K [NEB #P8107S] in 2× SSC) and incubated at 37°C overnight. Embedded and cleared samples were thoroughly washed in 2× SSC to remove residual detergent and stored at 4°C prior to imaging.

#### MERFISH and sequential FISH imaging

Two-color MERFISH was performed with a fully custom imaging setup as previously described^78^. Briefly, samples were stained at the bench with the first cycle of fluorescent readout probes (targeting *Oxt* mRNA and poly(A) anchors) in 2-3 mL wash buffer (10% ethylene carbonate (EC; Sigma #E26258), 1 mM EDTA, 0.1% Triton X-100 in 2× SSC) for at least 1 hour. Samples were stained with 4′,6-diamidino-2-phenylindole (DAPI) in wash buffer for 10 minutes, then washed thoroughly with 2× SSC and trimmed to size. On the microscope, a 10X objective was used to collect DAPI mosaic for sample orientation. Approximately 9 x 11 fields of view (FOVs; 1.8 x 2.2 mm) were selected per tissue section for sequential FISH and MERFISH. Images were acquired using a Nikon Plan Apo 60X objective. For MERFISH, poly(A) and DAPI imaging, 6 z-slices were captured with 1.8 µm spacing. For sequential FISH, 4 z-slices were captured with 3 µm spacing.

A typical imaging and fluidics cycle involves the following steps. (i) The sample chamber is perfused with imaging buffer (0.5 mg/mL Trolox, 50 µM Trolox-Quinone, 5 mM 3,4dihydroxybenzoic acid, 0.2% v/v recombinant bacterial procatechuate oxidase (OYCAmericas, 46852004), 5 mM EDTA in 50 mM Tris-HCl pH 8.0, adjusted to pH ∼8.8 with 5 mM NaOH), and allowed to equilibrate for at least 1 minute. (ii) The sample is imaged with up to five laser lines (750, 635, 545, 473, 405 nm). (iii) Fluorophores are removed by flowing 50 mM tris-2(-carboxyethyl)phosphine (TCEP; Cat) in 2× SSC for 15 minutes. (iv) The sample chamber is washed with 2× SSC. (v) Samples are hybridized with 3 nM fluorescent readout probe in EC wash buffer. (vi) The sample chamber is washed with EC wash buffer to remove residual unbound probes.

#### MERFISH and sequential FISH image processing

Images were registered to the first cycle of imaging (containing *Oxt* mRNA, polyA anchor and DAPI) using fiducial beads and decoded using a previously described pipeline (github.com/ZhuangLab/MERFISH_analysis)^50,74^. Due to occasional large microscope stage shifts between imaging separately libraries, on the order of 10-20 µm, individual library runs were further registered to the first cycle via phase correlation on the Laplacian-transformed fiducial bead images. For segmentation, poly(A) and DAPI images were downsampled to 512 x 512 pixels and stitched into mosaics. Illumination characteristics of the field were estimated on a run-to-run basis using the basicpy package, and the resultant flatfield correction was applied to each image. Overlapping image edges were further smoothed by linear interpolation to obtain a contiguous mosaic. Cell boundaries were predicted using CellposeSAM^79^ on the complete stitched mosaics, and transcripts were assigned to the corresponding masks. For sequential FISH, intensity per cell volume was assigned to the corresponding masks. The final RNA identities, associated metadata for decoding, and cell assignments are provided at Dryad (10.5061/dryad.v6wwpzh98).

#### Single-cell analysis and label transfer

Single cell analyses were carried out using scanpy^80^. Spatial autocorrelation analyses were performed using squidpy. To obtain a common embedding between scRNA-seq data from the Allen Brain Institute and previously published snRNA-seq from the P65 hypothalamus and our MERFISH data^49,53^, we leveraged a deep variational autoencoder method, scVI^81^, which represents the transcriptome of a given cell in a given number of latent dimensions while correcting for batch effects. In addition, a second scVI model was used to embed the full MERFISH dataset only.

Label transfer was achieved using a two-step process with scANVI. First, high-level labels were transferred from the Allen Brain atlas to call non-neuronal and neuronal populations. The Allen Brain terminal labels were applied to the non-neuronal cells, and neuronal populations were subsetted for the second round of label transfer. In this second round, we reran scANVI to transfer high-resolution hypothalamic labels from our previous P65 hypothalamus atlas, where we previously provided approximate regional localizations and associations to behavioral phenotypes based on a survey of the literature. Due to regional differences between the snRNA and MERFISH datasets, for each P65 cell type, we manually checked for spatially coherent patterns and observed patterns mostly consistent without our previous annotation set. A subset of cell types were excluded because they had few cells across our datasets and, thus, were too infrequently observed for robust measures of expression. In addition, we removed slices and cell populations assigned to slices that were imaged with MERFISH but which fell outside of the region characterized with in this published dataset. All single cell adata objects are provided as-is, with an additional flag *ignoreCells* used to exclude those cells from downstream analyses. MERFISH single-cell data are provided at Dryad (10.5061/dryad.v6wwpzh98).

Differential gene expression was performed using pydeseq2^72^ on pseudobulk transcript counts on a per cell-type basis. Gene set enrichment analysis on the resultant shrunken log2-fold change estimates was performed using gseapy^82^.

---

**Figure S1.**
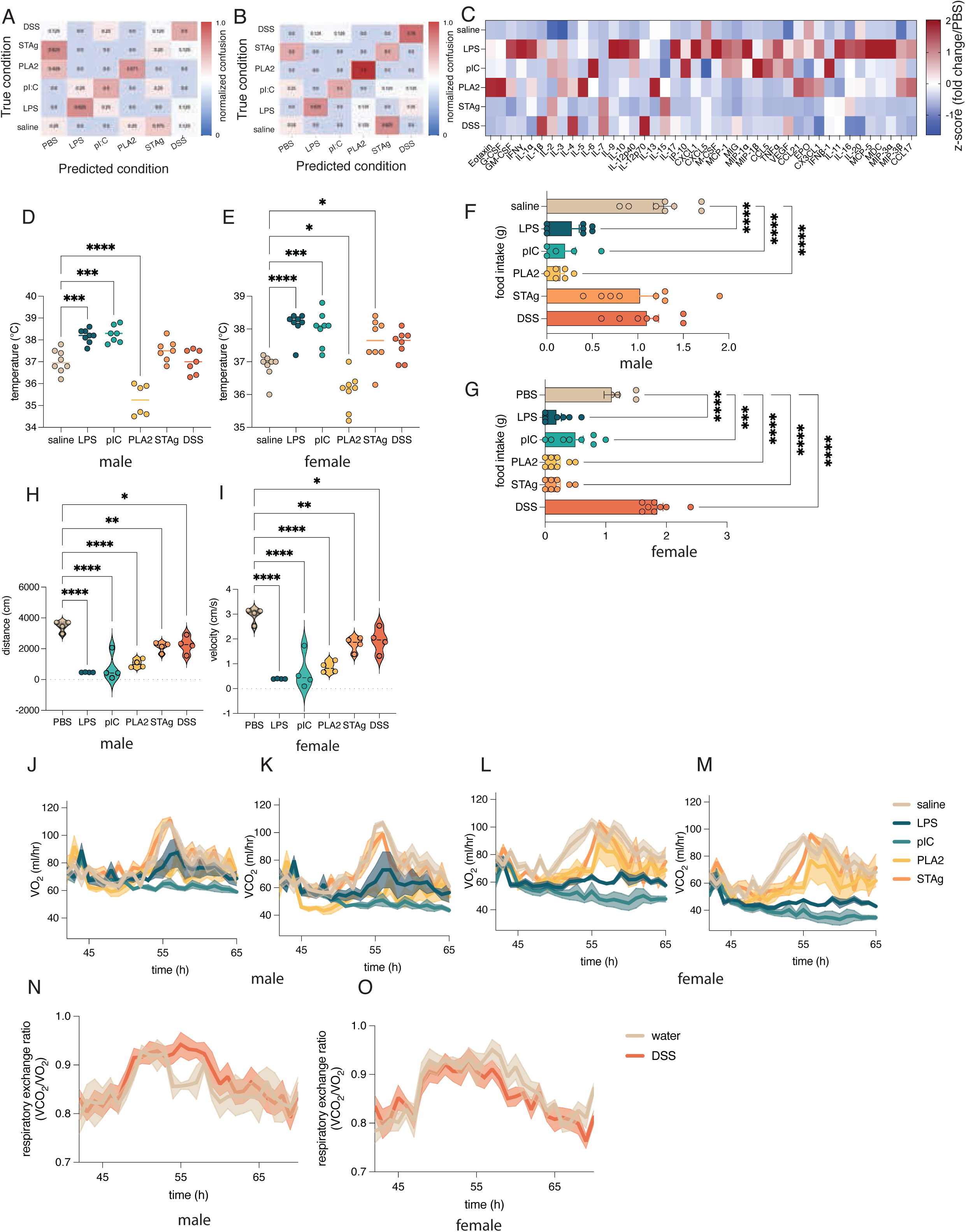
Related to Figure 1. (A) Normalized confusion matrices showing accuracy of random forest classifier for predicting sickness state based on 42 measured serum cytokines. F1 = 0.525 +/- 0.085. (B) Normalized confusion matrices showing accuracy of random forest classifier for predicting sickness state based on serum cytokines shown in Fig. 1B F1= 0.597 +/- 0.158. (C) z-scored concentration of 42 serum cytokines measured by multiplexed ELISA in males and females during different sickness states. (D) Temperature in male FVB mice during different sickness states (E) Body temperature in female FVB mice during different sickness states (F) Food intake in male FVB mice during different sickness states over 4h (G) Food intake in FVB female mice during different sickness states over 4h (H) Distance traveled by male FVB mice during different sickness states over 10 minutes (I) Average locomotor velocity of female FVB mice during different sickness states over 10 minutes (J) VO_2_ in B6 male mice during different sickness states (K) VCO_2_ in B6 male mice during different acute sickness states (L) VO_2_ in B6 female mice during different acute sickness states (M) VCO_2_ in B6 female mice during different sickness states (N) RER in B6 male mice during DSS challenge (O) RER in B6 female mice during DSS challenge. p<0.05* p<0.01** p<0.001*** p<0.0001**** based on ordinary one-way ANOVA with Dunnett’s multiple comparisons test. All measurements taken at 4h postinjection (acute models) or day 3 (DSS).

**Figure S2.**
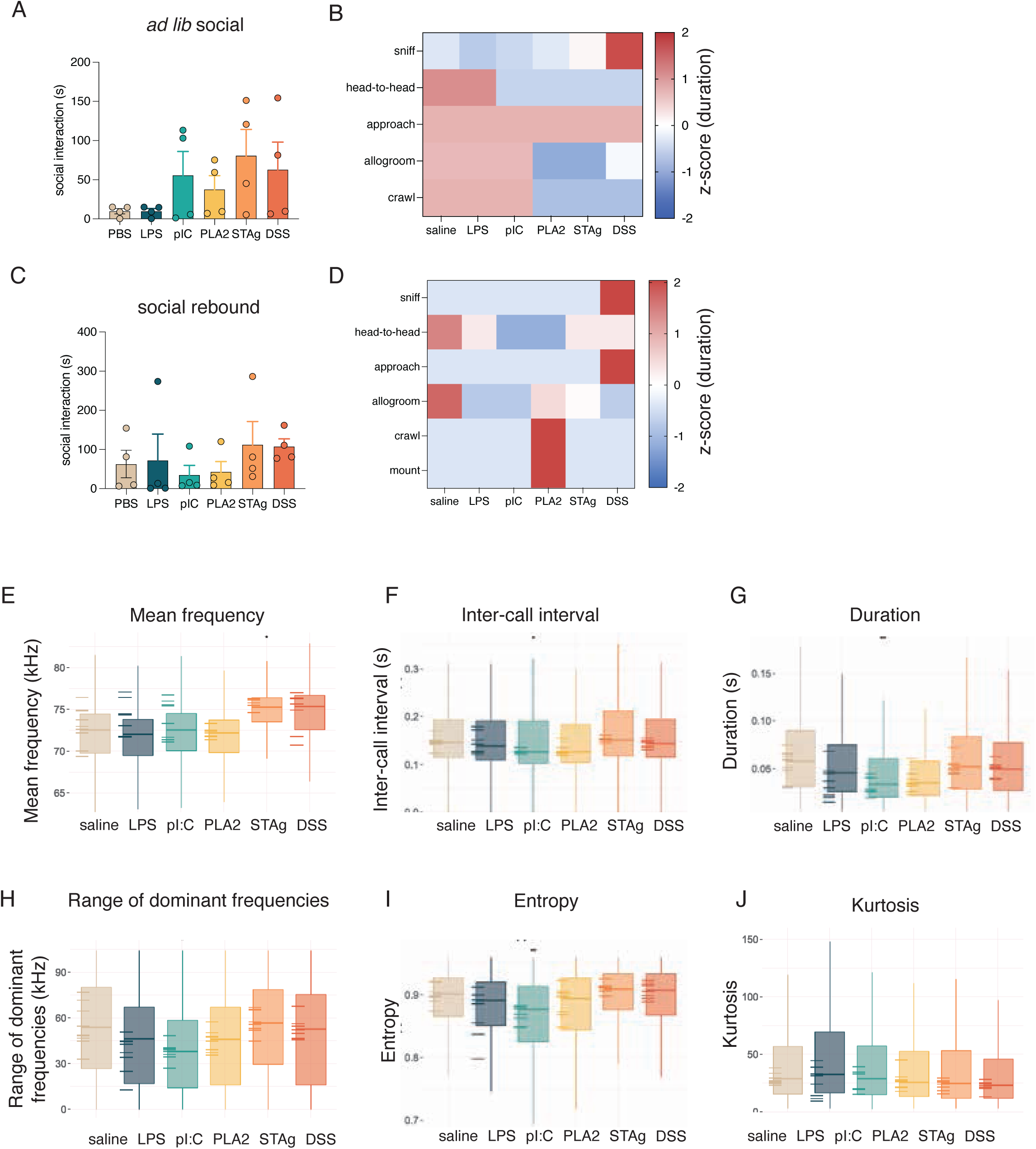
Related to Fig. 2 (A) (*Ad libitum* social interaction in C57BL/6 female mice recorded over 10 minutes.B) summary of individual social behaviors quantified in (A). (C) social rebound in B6 female mice recorded over 10 minutes. (D) summary of individual behaviors quantified in (C). (E-J) subset of USV acoustic features computed using WarbleR Heat maps represent zscored data. All measurements taken at 4h post-injection (acute models) or day 3 (DSS).

**Figure S3.**
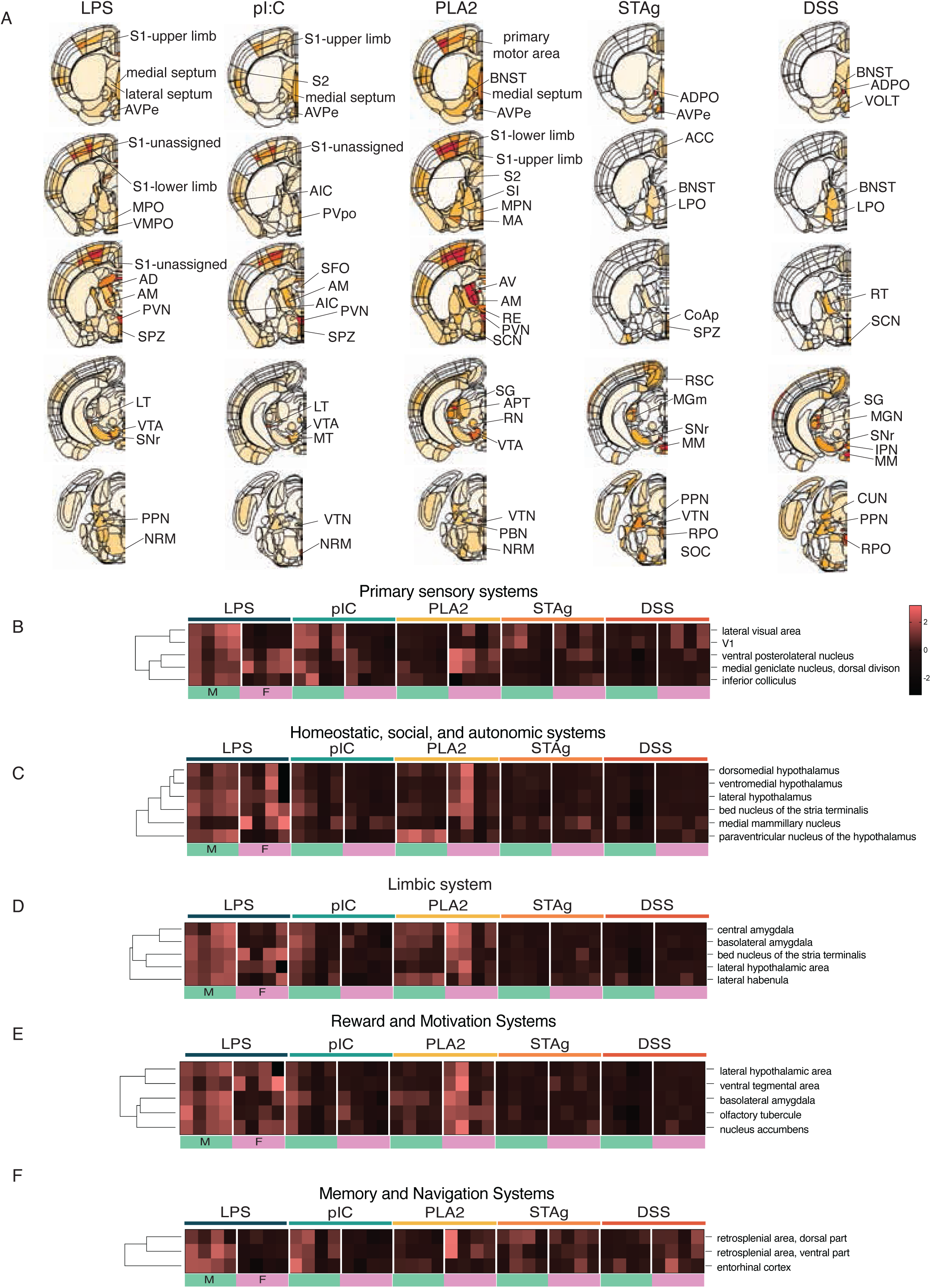
Related to Fig. 3 (A) Graphical heat map of average FOS intensity across coronal sections in all sickness states, normalized to saline controls. (B-F) Hierarchical clustering of FOS intensity normalized to saline controls in brain areas grouped by functional categories, separated by sex. Acronyms of brain areas shown in (A) listed in Table S1.

**Figure S4.**
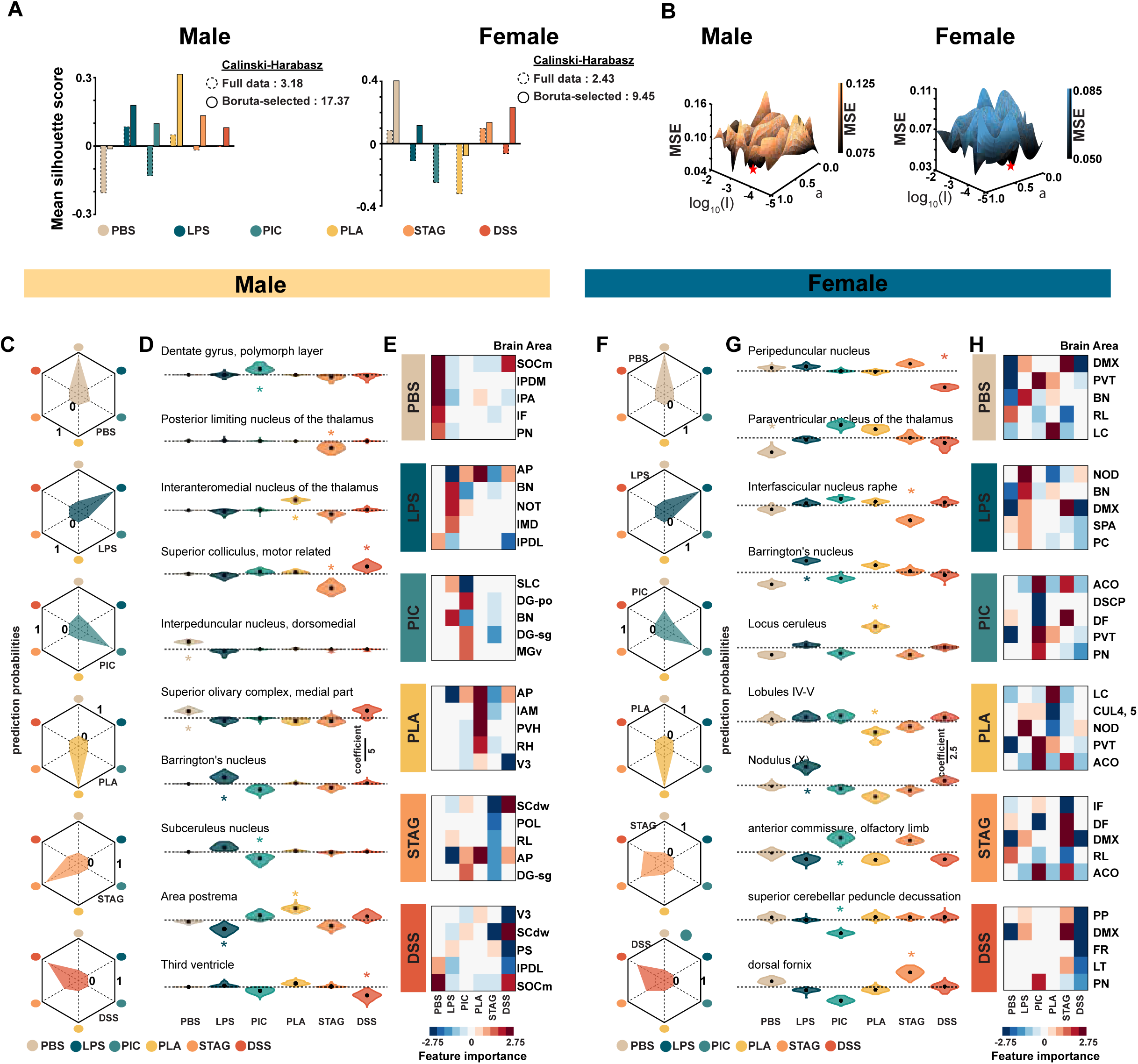
Related to Fig. 4: Validation of feature selection and model stability. (A) Quantitative validation of clustering enhancement: Mean silhouette score and Calinski-Harabasz index for full (641 areas) and Boruta-selected (36/25 areas) data. The Boruta feature selection significantly improved data partition. The silhouette distance became more positive (or less negative) in both sexes (left: males, right: females) after selection. The Calinski-Harabasz index showed a dramatic increase from 3.18 to 17.37 in males and 2.43 to 9.45 in females. (B) Elastic net hyperparameter selection: plots illustrating the Mean squared error (MSE) dependence on the Elastic net regularization hyperparameters, λ and α, used during model training. The choice of λ and α was optimized for robustness in handling highdimensional, potentially correlated FOS data. Prediction probabilities and feature importance in males (C-E) and females (F-H): Prediction probabilities for individual test samples across the six sickness states in males (C) and females (F), showing high prediction probabilities were consistently assigned to the true sickness state. Plots showing the distribution of coefficients across all cross-validation iterations and sickness states, providing insight into the stability and consistency of each region’s predictive influence in males (D) and females (G). Heatmaps depicting a hybrid metric, defined as | SHAP|*β, combining the magnitude of SHAP-style contributions with the stability of the model coefficients for males (E) and females (H).

**Figure S5.**
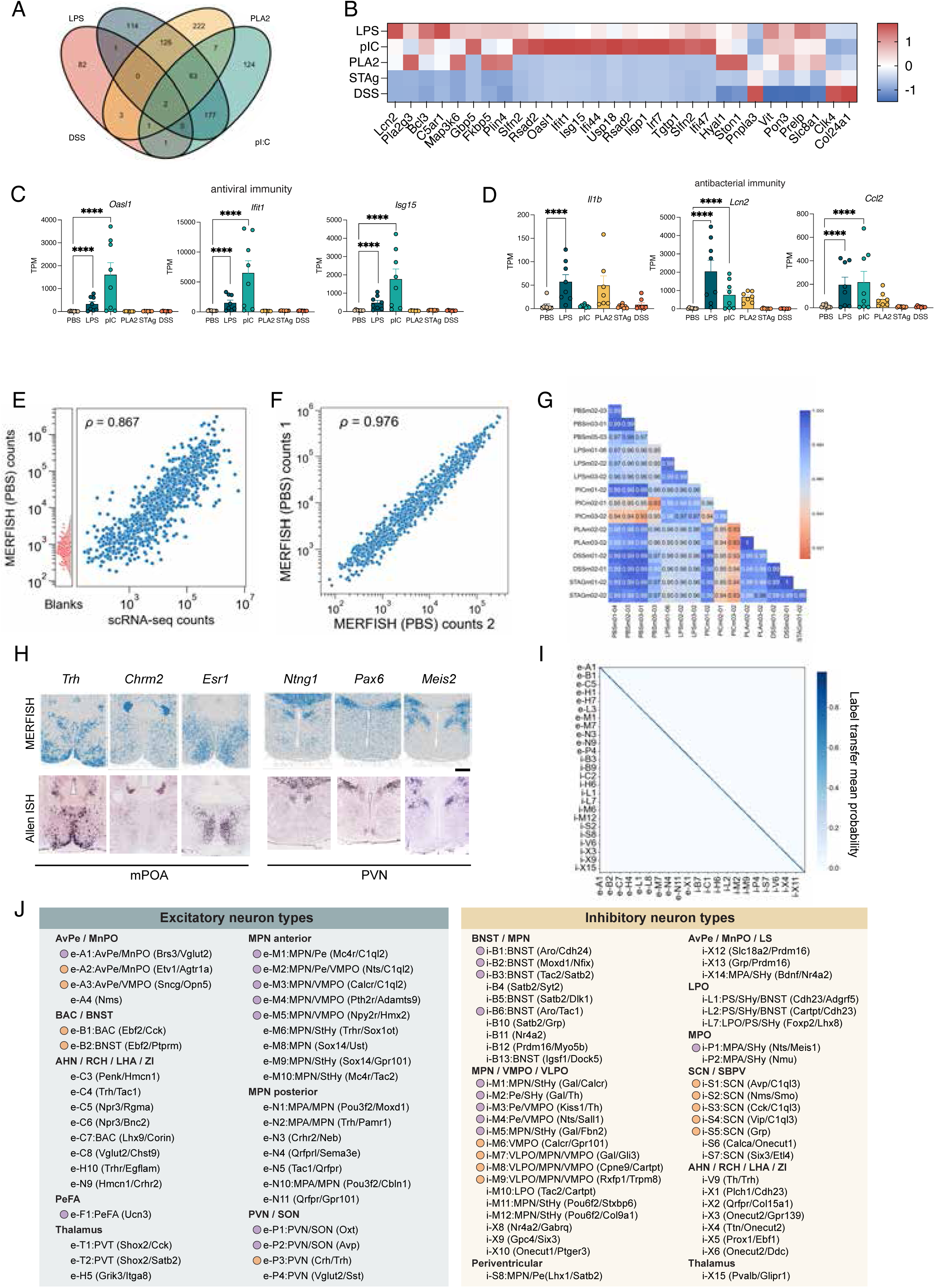
Related to Fig. 5 (A) Number of differentially-expressed genes identified by bulk RNAseq in the POA and PVN across conditions. (B) Top DEGs in each condition assessed by bulk RNAseq (C&D) expression genes of interest assessed by bulk RNAseq. (E) Correlation between MERFISH and scRNA-seq counts from the Allen Brain Institute (hypothalamic region). (F) Correlation between MERFISH counts from two PBS-treated animals. (G) MERFISH sample-to-sample correlation. (H) Example spatial localizations of MERFISH genes (above) and corresponding Allen Brain in situ hybridization (ISH) data in the mPOA and PVN. (I) Label transfer from P65 hypothalamus Multiome snRNA. Heatmap shows mean label probability applied to MERFISH data. Only a subset of labels are shown. (J) Summary of all neuronal cell-types annotated using MERFISH and correspondence to known functions. Orange circles indicate cell-types involved in homeostatic behaviors. Purple circles represent cell-types involved in social behavior. Scale = 500µm.

**Figure S6.**
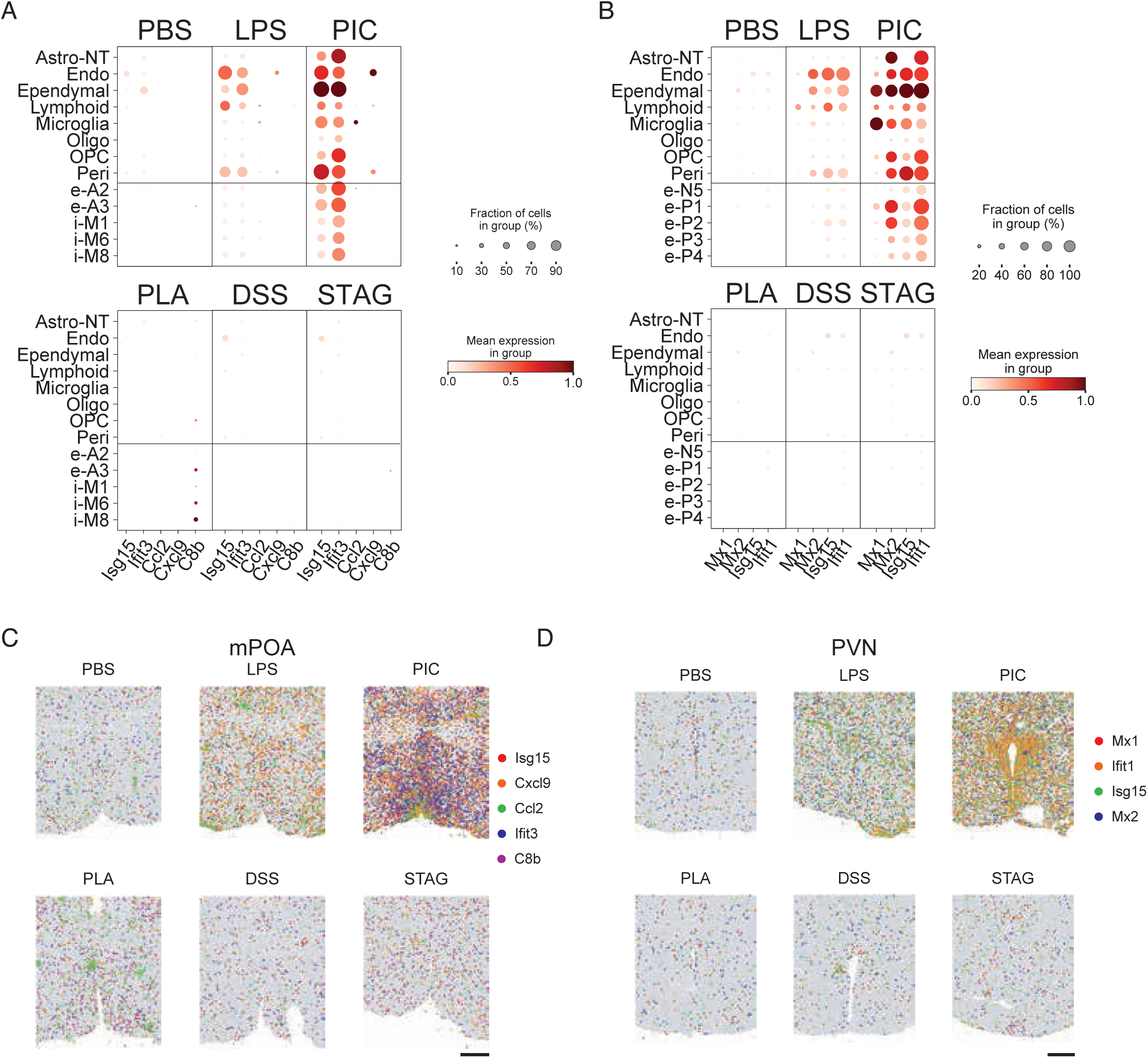
Related to Fig. 6 (A) Cell type-specific gene expression changes in the POA (B) Cell type-specific gene expression changes in the PVN (C) Spatial location of upregulated immune genes in the POA (D) Spatial location of upregulated immune genes in the PVN. Scale = 500µm

## Notes

https://github.com/Dulac-Lab/Brain-immune-interactions

### Competing Interest Statement

J.R.M. is an inventor of patents applied for by Harvard University and Boston Childrens Hospital related to MERFISH. J.R.M. is a co-founder and consultant of Vizgen, Inc. The other authors declare no conflict of interest.

